# INPP5D/SHIP1 regulates endo-lysosomal function and cargo-selective phagocytosis in human microglia

**DOI:** 10.1101/2025.10.27.684632

**Authors:** Gizem Terzioglu, Emma S. Karp, Sarah E. Heuer, Jean Paul Chadarevian, Fasih M. Ahsan, Verena C. Haage, Hayk Davtyan, Garrett M. Fogo, Annabel J. Curle, Nancy Ashour, Courtney R. Benoit, Jillian K. Schmottlach, Anna E. Cramer, Christina R. Muratore, Duc M. Duong, David A. Bennett, Nicholas T. Seyfried, Philip L. De Jager, Mathew Blurton-Jones, Tracy L. Young-Pearse

**Affiliations:** Ann Romney Center for Neurologic Diseases, Department of Neurology, Brigham and Women’s Hospital and Harvard Medical School, Boston, MA, USA; Department of Neurobiology & Behavior, Institute for Memory Impairments and Neurological Disorders and Sue and Bill Gross Stem Cell Research Center, University of California, Irvine, Irvine, CA, USA; Center for Translational & Computational Neuroimmunology, Department of Neurology and the Taub Institute for Research on Alzheimer’s disease and the Aging Brain, Columbia University Irving Medical Center, New York, NY, USA; Department of Biochemistry, Emory School of Medicine, Atlanta, GA, USA; Rush Alzheimer’s Disease Center, Rush University Medical Center, Chicago, IL, USA

## Abstract

*INPP5D*, which encodes SHIP1, is genetically associated with Alzheimer’s disease (AD) risk and regulates microglial immune function. Here we identified SHIP1 as a regulator of endo-lysosomal homeostasis in human microglia. *INPP5D* haploinsufficiency impaired endosome maturation and lysosomal degradation, leading to lipid droplet accumulation, lysosomal stress, and NLRP3 inflammasome activation. SHIP1-deficient microglia shifted from an immune-responsive state toward a disease-associated state with elevated cargo-selective phagocytosis. Although amyloid-β uptake was unaffected, SHIP1-deficient microglia accumulated intracellular amyloid-β and exhibited exacerbated lipid droplet accumulation, consistent with defective lysosomal processing. Proteomic analysis of genetically diverse human microglia revealed that the protective *INPP5D* rs10933431 variant is associated with elevated phosphatase-domain containing SHIP1, and protein profiles consistent with enhanced endolysosomal trafficking. In mice xenografted with SHIP1-deficient human microglia, high amyloid-β burden exacerbated lysosomal, lipid droplet and inflammasome phenotypes. Together, these findings identify disrupted endo-lysosomal trafficking and degradative capacity as mechanisms linking *INPP5D* dysfunction to disease-associated microglial states in AD.

## INTRODUCTION

Src homology 2 (SH2) domain containing inositol polyphosphate 5-phosphatase 1 (SHIP1) is a membrane-associated lipid phosphatase that removes the 5’ phosphate from the second messenger PI(3,4,5)P_3_ to generate PI(3,4)P_2_, thereby negatively regulating PI3K/AKT signaling. In addition to its phosphatase activity, SHIP1 functions as a scaffolding protein through its additional domains, enabling interactions with proteins containing immunoreceptor tyrosine inhibitory or activating motifs (ITIMs/ITAMs), as well as SH2 and SH3 domains^1^. Through the integration of its enzymatic and scaffolding functions, SHIP1 has emerged as a key negative regulator of inflammation across multiple myeloid and lymphoid signaling pathways^2^. As such, dysfunction of SHIP1 has been associated with a spectrum of diseases, including Alzheimer’s disease (AD), acute myeloid leukemia (AML) and autoimmune diseases such as systemic lupus erythematosus (SLE) and rheumatoid arthritis (RA)^3–6^. Therefore, elucidating the biology of SHIP1 and the molecular outcomes of its dysfunction is essential for uncovering how SHIP1 shapes immune responses and contributes to diverse pathologies.

Microglia, the resident immune cells of the central nervous system (CNS), play a variety of roles in the pathogenesis of late-onset AD, a disease involving extensive neuroinflammation and neurodegeneration. Recent genome-wide association studies (GWAS) have identified several genes associated with AD that are predominantly or exclusively expressed in myeloid cells, including *INPP5D* (inositol polyphosphate 5-phosphatase 1), which encodes SHIP1^7–10^. Single nucleotide polymorphisms (SNPs) at the *INPP5D* locus have been identified that are associated with AD. These SNPs are intronic and it is unclear how they influence *INPP5D* levels^11,12^. Intriguingly, RNA expression data from human brain tissue reveal the presence of isoforms of *INPP5D* that encode truncated SHIP1 that lacks phosphatase domain^12^. In accord, while overall protein levels of SHIP1 are elevated in brain tissue from individuals with AD, the levels of full length SHIP1 containing the phosphatase domain are reduced^11^. Thus, both genetic and neuropathological evidence links *INPP5D* to AD, underscoring the need to define how SHIP1 regulates microglial biology and how this may contribute to disease pathogenesis.

In line with SHIP1’s role as a brake on inflammation, we recently identified SHIP1 as a regulator of the NLRP3 inflammasome in human microglia, as both the CRISPR/Cas9-mediated knockdown of *INPP5D* and acute inhibition of SHIP1 led to the activation of the inflammasome^11^. NLRP3 inflammasome is a multiprotein complex that leads to the secretion of the pro-inflammatory cytokines IL-1β and IL-18 upon its activation, and its aberrant activation has been extensively linked to AD^13^. Canonically, inflammasome activation requires two stimuli: an inflammatory “priming” signal that induces NF-κB–dependent transcription of inflammasome components, followed by a second inflammatory stimulus that triggers the assembly of the NLRP3 inflammasome^14^. However, recent studies show that dysfunction of organelles, such as lysosomes and mitochondria, can activate the NLRP3 inflammasome in the absence of external stimuli^15^. Thus, a key question is how microglial SHIP1 deficiency drives inflammasome activation in the absence of external stimuli, raising the possibility that organelle dysfunction provides the endogenous cue for NLRP3 activation.

Here, using human induced pluripotent stem cell (iPSC)-derived microglia (iMGs), we identify SHIP1 as a regulator of endo-lysosomal function. We show that SHIP1 localizes in part to endo-lysosomal compartments and binds to the CapZ family of proteins, which are important for endosome maturation. We find that reduction of SHIP1 levels via CRIPSR/Cas9-mediated genome editing in iMGs impairs lysosomal homeostasis and degradation, leading to lipid droplet accumulation and leakage of lysosomal cathepsin B into the cytosol, which mediates the activation of the NLRP3 inflammasome. Moreover, we identify a shift from an immune-responsive state to a disease-associated phagocytic state in microglia with loss of one copy of *INPP5D*, which renders these microglia insensitive to inflammatory stimulus while hyper-phagocytic towards synaptic material and apoptotic neurons. We further show that antibody-mediated modulation of TREM2 partially rescues excessive phagocytosis and impaired lysosomal degradation in SHIP1-deficient microglia. Despite no change in amyloid-β uptake, SHIP1 deficiency in microglia leads to intracellular accumulation of amyloid-β due to impaired degradative processing and further exacerbates lipid droplet accumulation. Finally, proteomic analysis of genetically diverse human microglia revealed that reduced SHIP1 abundance and the AD-associated rs10933431 risk allele are linked to altered endo-lysosomal programs, while brain-wide engrafted SHIP1-deficient human microglia in xenotolerant, amyloid accumulating 5xFAD-hFIRE (5x-hFIRE) mice exhibited exacerbated lysosomal, lipid droplet and phenotypes consistent with inflammasome activation. Together, these findings implicate loss of SHIP1 in microglia as a central driver of multiple pathogenic phenotypes associated with Alzheimer’s disease.

## RESULTS

### SHIP1 interacts with proteins important for endo-lysosomal function

To gain insight into the roles of SHIP1 in microglia, we investigated the interactome of SHIP1 in human microglia by performing immunoprecipitation followed by mass spectrometry (IP-MS) using iPSC-derived microglia (iMGs) derived from two different induced pluripotent stem cell (iPSC) lines from the Religious Orders Study and Rush Memory and Aging Project (ROSMAP) (Fig. 1a)^16^. One donor was a male (BR33), while the other was a female (BR24), both were over 90 years old at the time of their death with no cognitive impairment or other neurological diagnoses^17^. We generated an *INPP5D* knockout (KO) line using CRISPR/Cas9 in the BR24 cell line to use as a negative control for SHIP1 co-immunoprecipitation (co-IP) analyses. We successfully immunoprecipitated endogenous SHIP1 in both wild-type (WT) lines, with no detectable SHIP1 remaining in the post-IP supernatant (flowthrough) (Fig. 1a) and detected the proteins that co-immunoprecipitated with SHIP1 via mass spectrometry (Supplementary Table 1). We identified 38 proteins that were significantly (adj. p-value < 0.05) enriched across both genetic backgrounds more than 1.5-fold in WT microglia compared to KO microglia (Figs. 1c-e). We detected several SHIP1 binding partners previously identified in other myeloid cells, such as SH3KBP1^18^, GRB2^19^, SHC1^19^, DOK2^19,20^, ARAP1^21^ and CD2AP^22^ (Fig. 1d). Remarkably, SHIP2/INPPL1, which is the paralog of SHIP1 also implicated in AD^23^, was the strongest hit detected in the IP-MS, which we validated via co-immunoprecipitation followed by Western blotting (WB) (Fig. 1f), suggesting that SHIP1 and SHIP2 may act in concert to perform phosphatase-dependent and -independent functions in microglia.

**Fig. 1:**
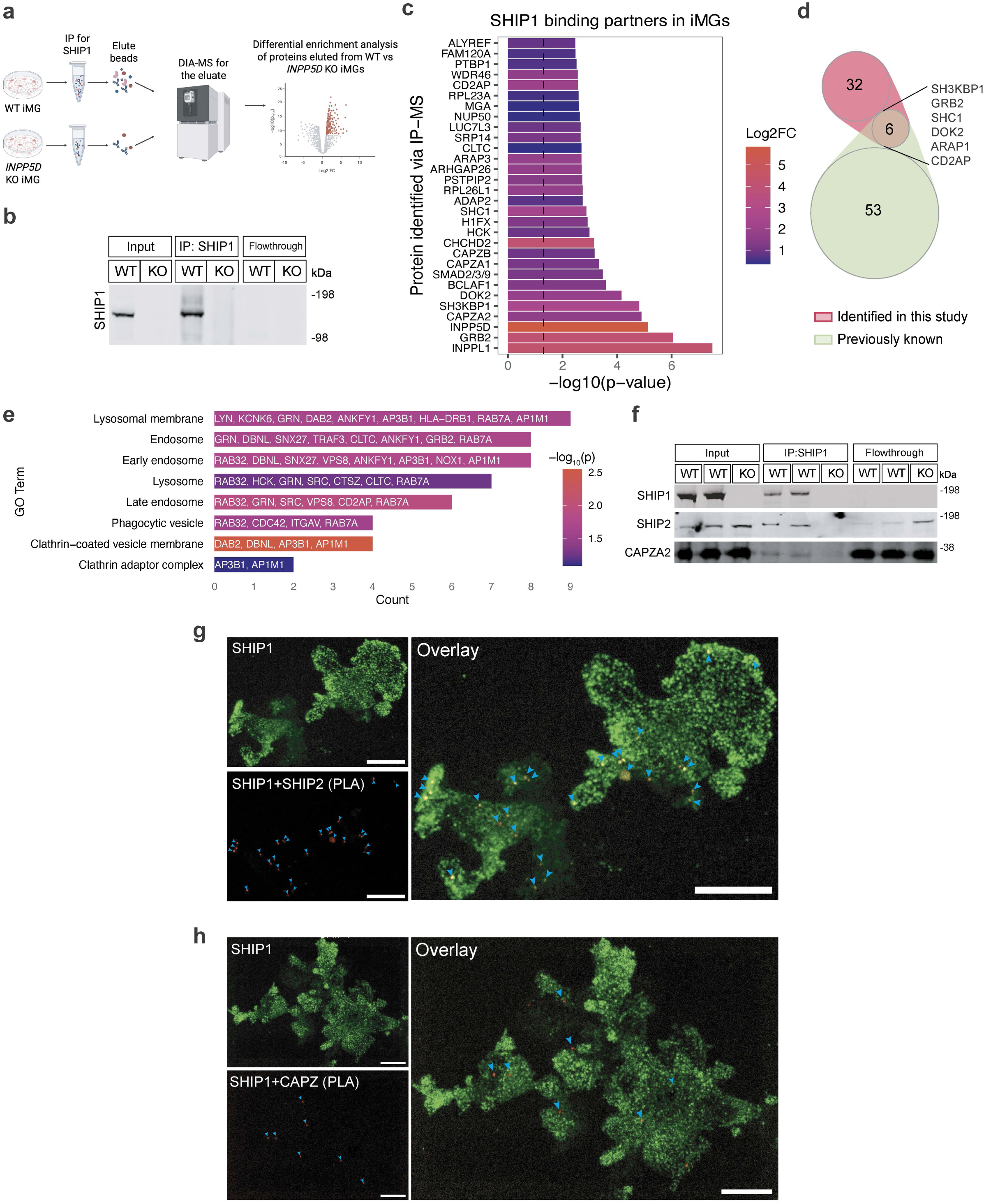
SHIP1 interacts with proteins important for endo-lysosomal function. a. Simplified experimental pipeline for SHIP1 immunoprecipitation followed by mass spectrometry (IP-MS). Created with BioRender.com. b. Representative Western blot of SHIP1 immunoprecipitation WT and *INPP5D* KO iMGs. c. Bar plot of the top 30 statistically significant binding partners of SHIP1 identified via IP-MS. d. Venn diagram of the previously identified SHIP1 binding partners and the binding partners identified in this study. e. SHIP1 binding partners identified via IP-MS categorized by endo-lysosomal pathway-related GO terms. f. Representative Western blot for the validation of co-immunoprecipitation of SHIP1 with SHIP2 and CAPZA2. g. Representative confocal image of SHIP1 immunostaining in BR24 iMGs, along with proximity ligation assay (PLA) signal showing areas of close interaction of SHIP1 and SHIP2. Scale bars: 25 μm. See Extended Data Fig. 1a for negative and positive controls. h. Representative confocal image of SHIP1 immunostaining in BR24 iMGs, along with proximity ligation assay (PLA) signal showing areas of close interaction of SHIP1 and CAPZ. Scale bars: 10 μm. See Extended Data Fig.1b for negative and positive controls.

The second strongest novel hit we identified was CAPZA2, a subunit of the F-actin capping protein complex (CapZ), which we validated by co-immunoprecipitation and WB (Figs. 1c,f). Of note, two other subunits of this protein complex, CAPZA1 and CAPZB, also co-immunoprecipitated with SHIP1 (Fig. 1c). The CapZ complex recently has been shown to modulate endosomal trafficking and early endosome maturation by regulating the F-actin density around early endosomes and facilitating Rab5 activation, implicating a potential role for SHIP1 in the endo-lysosomal system via its interaction with the CapZ complex^24,25^.

We used proximity ligation assay (PLA) to further validate SHIP1’s interaction with SHIP2 and CapZ in microglia *in situ* (Figs. 1g-h). Robust PLA signal for both SHIP1-SHIP2 (Fig. 1g) and SHIP1-CapZ (Fig. 1h) pairs were detected, indicating close spatial association of SHIP1 with both proteins in human microglia. In contrast, no signal was observed in INPP5D KO iMGs or in negative control conditions lacking the primary or secondary antibodies, confirming assay specificity (Extended Data Fig. 1).

SHIP1 is known as a cytoplasmic protein that is recruited to the plasma membrane upon immune receptor activation, where it hydrolyzes its lipid substrate PI(3,4,5)P_3_ into PI(3,4)P_2_^26^. Accordingly, the majority of the SHIP1 binding partners we identified via IP-MS are also cytoplasmic or membrane-associated proteins (Supplementary Table 2). Functionally, many of SHIP1’s binding partners are important for cytokine signaling, cellular response to stress, and antigen presentation (Supplementary Table 3), in line with SHIP1’s known roles in regulating immune response in myeloid cells^2^. Another pathway that SHIP1’s binding partners were significantly enriched in was “vesicle-mediated transport” (Supplementary Table 3). Further investigation of the binding partners revealed unexpected proteins that have known functions in various components along the endocytic pathway, including endosomes and lysosomes, hinting that SHIP1 may play a role in endo-lysosomal function through its interactions with these proteins (Fig. 1e). This led us to interrogate the integrity of the endo-lysosomal system in human microglia with SHIP1 deficiency.

### SHIP1 regulates endo-lysosomal function

Next, we asked whether reduction of functional SHIP1 levels in microglia, as observed in AD^11^, affects endo-lysosomal function (Fig. 2a). We generated iMGs heterozygous for *INPP5D* (referred to as “HET”) using CRISPR/Cas9 in both BR24 and BR33 cell lines (Extended Data Fig. 2a-b). First, endocytosis was measured by treating iMGs with bovine serum albumin (BSA)-Alexa 488 for 1 hour in the presence or absence of the actin polymerization inhibitor cytochalasin D (cyto D). BSA fluorescence was lower in the presence of cyto D but was not different between WT and HET iMGs after the 1-hour incubation (Fig. 2b-c), suggesting no change in endocytosis of BSA in HET iMGs. Next, we assessed lysosomal degradation in iMGs using Dye-Quenched (DQ)-Red BSA, whose fluorescence is unquenched upon degradation by lysosomal proteases. DQ-BSA intensity was lower in HET compared to WT iMGs in both genetic backgrounds, with a greater effect size in BR24, and abolished in the presence of the V-ATPase inhibitor bafilomycin A1 (Fig. 2c). Taken together, these results suggest that SHIP1 deficiency does not affect endocytic uptake but impairs the degradation of endocytosed cargo in microglia.

**Fig. 2:**
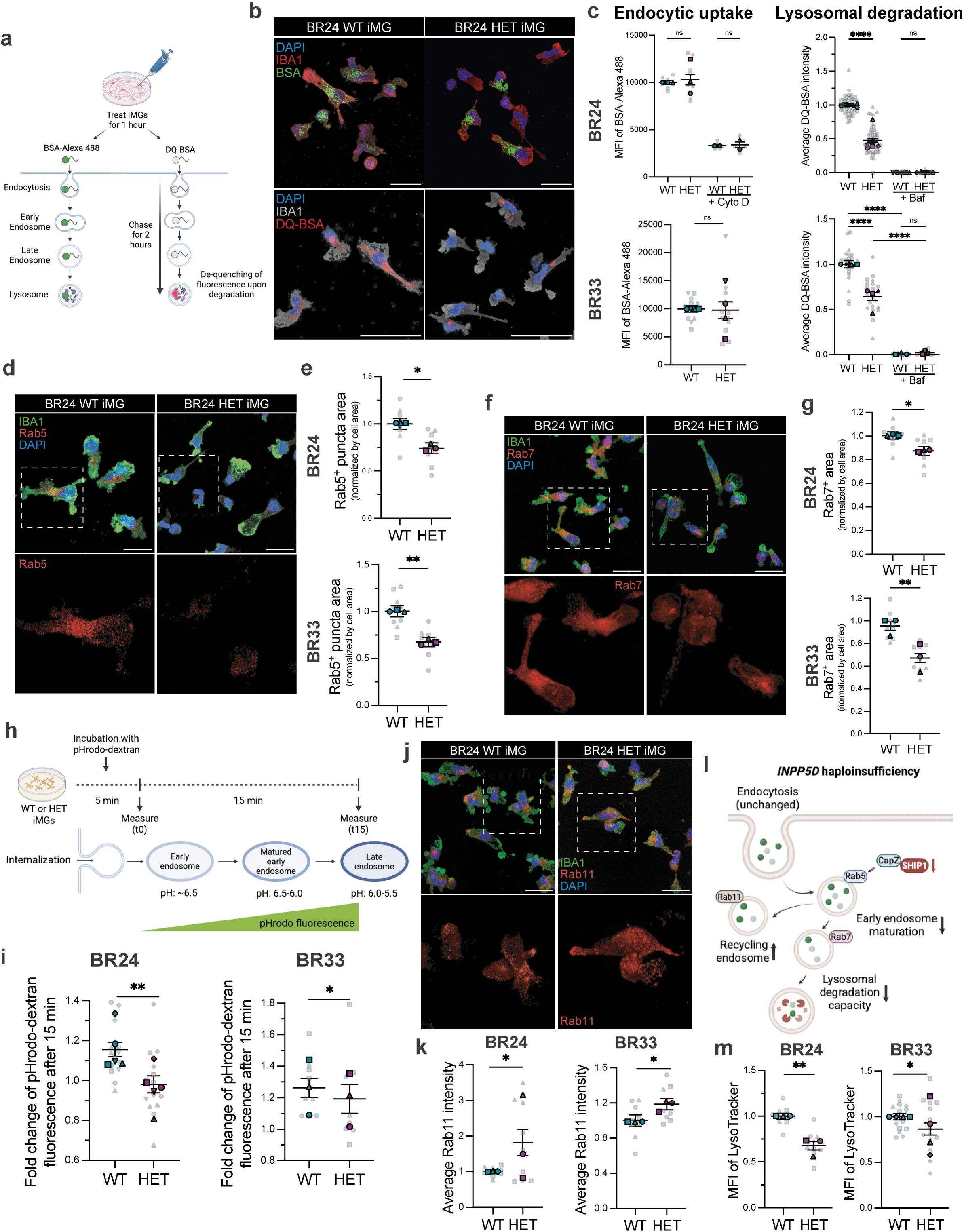
***INPP5D* haploinsufficiency in iMGs leads to endo-lysosomal dysfunction**. a. Schematic of the endocytosis and lysosomal degradation assays. Created with BioRender.com. b. Representative confocal images of BSA-Alexa488 and DQ-Red BSA within BR24 iMGs. Scale bars: 25 μm. c. Flow cytometric quantification of medium fluorescence intensity (MFI) of BSA-Alexa 488 and IncuCyte-based quantification of average DQ-Red BSA intensity within live BR24 and BR33 iMGs. Quantification shown for n=3 independent differentiations with 3 technical replicates per differentiation for BSA-Alexa4 488 assay. Quantification shown for n=4 independent experiments with 4-8 technical replicates per differentiation, each technical replicate averaged from 9 images per well for DQ-Red BSA assay. Two-way ANOVA was performed using averages from each differentiation. d. Representative confocal images of Rab5 in BR24 iMGs. Scale bars: 25 μm. See Extended Data Fig. 2d for images from BR33 iMGs. e. Quantification of Rab5+ puncta area in BR24 and BR33 iMGs. f. Representative confocal images of Rab7 in BR24 iMGs. Scale bars: 25 μm. See Extended Data Fig. 2g for images from BR33 iMGs. g. Quantification of Rab7+ puncta area in BR24 and BR33 iMGs. h. Experimental strategy for measuring endosome acidification in iMGs. Created with BioRender.com. i. Fold change of pHrodo-dextran fluorescence after 15 minutes of chase (t15), relative to the initial fluorescence (t0), used as a proxy for endosome maturation in BR24 and BR33 iMGs. See Extended Data Fig. 2h for representative histograms and gating strategy for pHrodo. j. Representative confocal images of Rab11 in BR24 iMGs. Scale bars: 25 μm. See Extended Data Fig. 2j for images from BR33 iMGs. k. Quantification of Rab11 intensity in BR24 and BR33 iMGs. l. Proposed model for consequences of *INPP5D* haploinsufficiency in microglia on endo-lysosomal trafficking and lysosomal degradation. Created with BioRender.com. m. Quantification of medium fluorescence intensity (MFI) of LysoTracker Green in live BR24 and BR33 iMGs, measured via flow cytometry. Unless otherwise stated, data is shown as mean ± SEM, normalized to WT conditions, obtained from n=3 independent differentiations with 3 technical replicates per differentiation. Mixed-effects analysis was performed with genotype and differentiation as fixed factors. The reported p-value corresponds to the main effect of genotype, averaged across differentiations. ****p < 0.0001, ***p < 0.001, **p < 0.01, *p<0.05; ns, not significant. For all scatter plots, different shapes delineate data from separate differentiations, average values per differentiation are shown, with faded dots representing technical replicates (wells) within each differentiation.

Immunostaining of microglia following the DQ-BSA assay revealed co-localization of a subset of SHIP1 with DQ-BSA+ puncta in WT iMGs. While there was reduced overall protein levels of SHIP1 in HET iMGs, the percentage of SHIP1 puncta co-localized with DQ-BSA was significantly higher in HET iMGs (Extended Data Fig. 2c, Supplementary Videos 1-2). Therefore, SHIP1 may localize to lysosomes, where it regulates endo-lysosome function via interactions with endo-lysosomal proteins.

While reduction of *INPP5D* does not affect endocytic uptake (Fig. 2c), we investigated whether endocytic maturation or trafficking is altered. Several binding partners of SHIP1, such as CD2AP^22^ and CapZ proteins (Fig. 1c) have known roles in early endosome trafficking and maturation^24,27^. Immunostaining revealed a significant reduction in Rab5+ and Rab7+ puncta area, which are markers for early and late endosomes respectively, in HET iMGs (Figs. 2d-g). On the other hand, the area of EEA1+ puncta, another marker for early endosomes, was unaffected (Extended Data Figs. 2d-g). These data indicate that early endosome abundance is preserved in HET iMGs, but the Rab5-to-Rab7 transition may be disrupted, prompting the examination of endosome maturation more directly. To assess endosome maturation, we measured acidification along the endocytic pathway using dextran fused to a pH-sensitive dye (pHrodo-dextran; Fig. 2h)^28,29^. pHrodo-dextran was measured directly after 5 minutes of incubation and after 15 minutes post-incubation, as previous studies established that internalized cargo localizes to late endosomes ∼10 minutes after uptake^30–32^. While the pHrodo signal in WT iMGs increased within 15 minutes, the increase in fluorescence was stunted in HET iMGs (Fig. 2i, Extended Data Fig. 2h-i). These results suggest that SHIP1 deficiency compromises Rab5-mediated early endosomal maturation and acidification, potentially leading to impaired delivery of internalized cargo for lysosomal degradation. To determine whether impaired maturation is accompanied by altered endosomal routing, we next examined Rab11, a marker of recycling endosomes. In contrast to Rab5, a decrease in Rab11+ puncta was not observed; instead, Rab11 signal intensity was increased in HET iMGs (Fig. 2j-k, Extended Data Fig. 2j-k), suggesting that SHIP1 deficiency may promote increased recycling of cargo that fails to progress efficiently to lysosomal degradation (Fig. 2l).

Given SHIP1’s potential interaction with lysosomal proteins (Fig. 1e) and its increased co-localization with lysosomes in HET iMGs (Extended Data Fig. 2e), we next assessed lysosomal compartments using LysoTracker Green, a cell-permeable dye that labels acidic organelles. Flow cytometric analysis revealed reduced LysoTracker intensity in HET iMGs (Fig. 2m), indicating a diminished pool of LysoTracker+ acidic organelles. This finding prompted us to investigate whether lysosomal integrity and stability is altered in HET iMGs.

### SHIP1 deficiency compromises lysosomal integrity

While impaired endosome maturation may explain the impairment in lysosomal degradation in HET iMGs, we previously reported a reduction in autophagic flux upon *INPP5D* loss-of-function in iMGs^11^, suggesting broad lysosome dysfunction. An important target for autophagic degradation is lipid droplets (LDs), the accumulation of which is considered a marker of inflammation in microglia during aging and Alzheimer’s disease^33–36^. To quantify LDs, iMGs were stained with BODIPY 493/503, a dye that detects LDs consisting of neutral lipids^35,37^. HET iMGs accumulate more LDs at baseline (Extended Data Figs. 3a-b), suggesting that SHIP1 deficiency impairs the degradation of both intracellular and extracellular cargo.

To elucidate the mechanism underlying the impairment in lysosomal degradation upon *INPP5D* haploinsufficiency, we first investigated LAMP1 levels in iMGs. In BR24 iMGs, LAMP1 protein levels measured by Western blotting and the relative area of LAMP1+ puncta normalized to IBA1+ area were both reduced in HET cells. In contrast, LAMP1 protein levels and puncta area were unchanged in BR33 HET iMGs (Extended Data Figs. 3c-e).

To assess lysosomal acidification at the single-organelle level, iMGs were transduced with a lentiviral ratiometric lysosomal pH reporter (FIRE-pHLy^38^; Extended Data Fig. 3f). While the mTFP1/mCherry ratio was increased in BR24 HET iMGs, suggesting reduced acidification, no change was observed in BR33 HET iMGs (Extended Data Fig. 3g), suggesting that genetic background modulates the impact of SHIP1 deficiency on lysosome abundance and pH. Notably, these findings did not fully account for the reduced LysoTracker signal observed across HET iMGs, particularly in the absence of consistent changes in lysosomal acidification. We therefore hypothesized that lysosomal membrane integrity may be compromised and next assessed lysosomal membrane permeabilization (LMP) using galectin-3 immunostaining. Galectin-3, which is normally diffuse in the cytoplasm, is recruited to lysosomes upon lysosomal membrane permeabilization and is widely used as a sensitive marker of lysosomal damage^38^. As a positive control, 1-hour treatment of iMGs with lysosomal damage-inducing L-leucyl-L-leucine methyl ester (LLoMe) resulted in clear galectin-3 puncta colocalized with LAMP1. HET iMGs exhibited increased galectin-3 signal intensity and a 3–4-fold increase in the proportion of cells containing galectin-3–positive puncta, indicating membrane permeabilization in a subset of lysosomes upon SHIP1 deficiency (Figs. 3a-c, Extended Data Fig. 4a). The concordance across the two genetic backgrounds of this phenotype supports an effect of SHIP1 levels on lysosomal permeabilization.

**Fig. 3:**
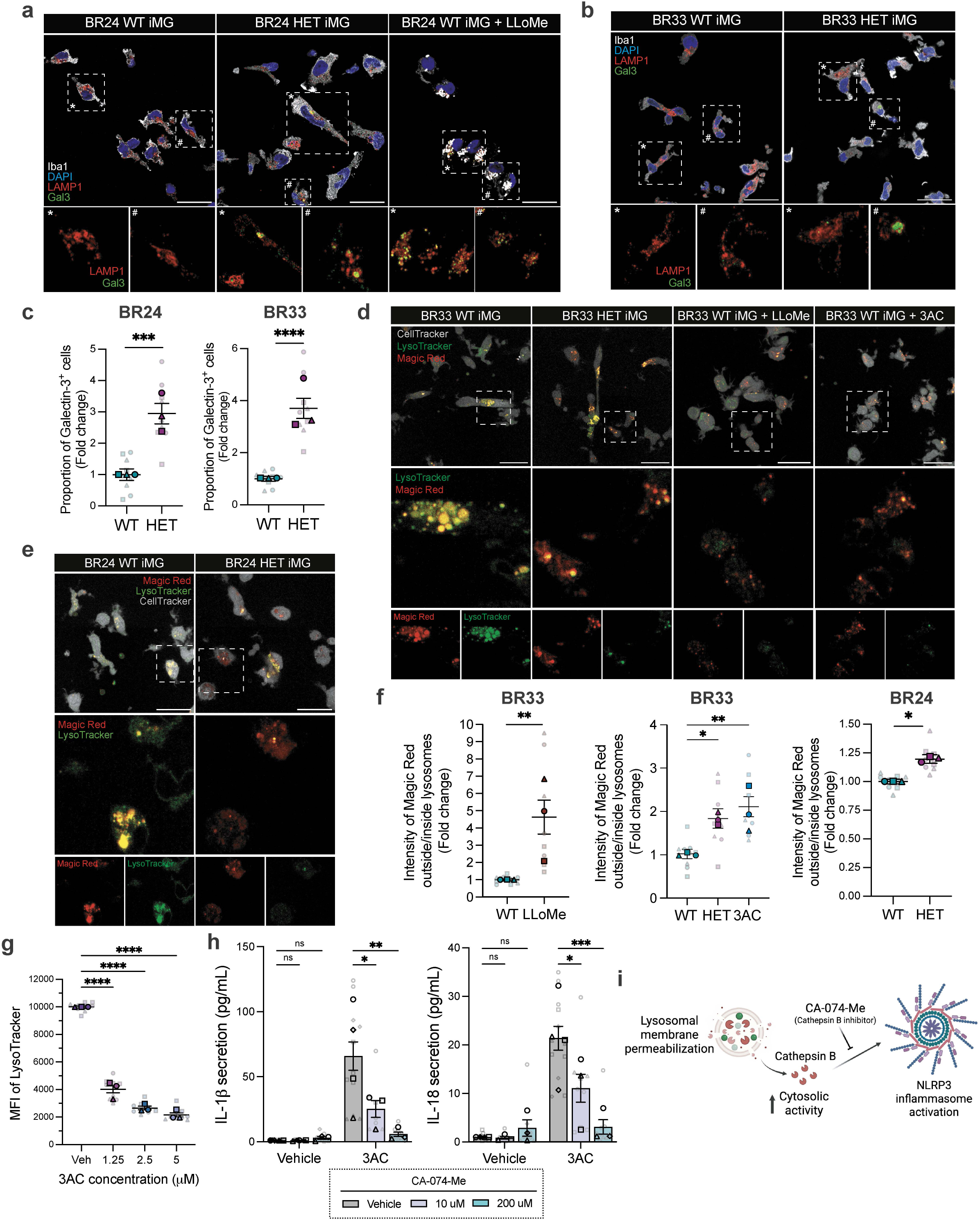
SHIP1 deficiency induces lysosomal dysfunction linked to NLRP3 inflammasome activation in human microglia. a. Representative confocal images of LAMP1 and galectin-3 in BR24 and (b) BR33 iMGs. LLoMe was used to induce lysosome damage as positive control. Scale bars: 25 μm c. Quantification of proportion of galectin-3+ puncta BR33 and BR24 WT and HET iMGs. d. Representative live confocal images of LysoTracker and Magic Red in BR24 and (e) BR33 iMGs. Scale bars: 25 μm f. Quantification of the ratio of Magic Red intensity outside versus inside LysoTracker+ lysosomes in BR24 and BR33 iMGs. g. MFI of LysoTracker in BR33 iMGs treated with different concentrations of 3AC for 6 hours. One-way ANOVA with Fisher’s least significant difference test was performed using averages from each differentiation. h. Quantification of IL-1β and IL-18 secretion, measured via ELISA (MSD) from iMGs treated with vehicle (EtOH) or 3AC for 6 hours and pre-treated with vehicle or different doses of CA-074-Me for 1 hour prior to the addition of vehicle/3AC. i. Proposed model for the mechanism of NLRP3 inflammasome activation downstream of SHIP1 deficiency or inhibition. Created with BioRender.com Unless otherwise stated, data is shown as mean ± SEM, normalized to WT conditions, obtained from n=3 independent differentiations with 3 technical replicates per differentiation. Mixed-effects analysis was performed with genotype and differentiation as fixed factors. The reported p-value corresponds to the main effect of genotype, averaged across differentiations. ****p < 0.0001, ***p < 0.001, **p < 0.01, *p<0.05; ns, not significant. For all scatter plots, different shapes delineate data from separate differentiations, average values per differentiation are shown, with faded dots representing technical replicates (wells) within each differentiation.

We previously reported that the NLRP3 inflammasome is activated in *INPP5D* HET microglia and in microglia following acute inhibition of SHIP1^11,15^. However, the mechanism underlying this inflammasome activation was unclear. Recently, it has become apparent that in some systems, lysosomal dysfunction may trigger inflammasome activation^15^. Specifically, lysosomal cathepsins, such as cathepsin B, can induce NLRP3 inflammasome activation upon leaking into the cytosol following LMP^39,40^. We therefore hypothesized that cathepsin B might mediate the NLRP3 inflammasome activation downstream of SHIP1 deficiency. Cathepsin B activity was measured using Magic Red, a fluorogenic substrate that fluoresces upon cleavage by cathepsin B. Cathepsin B activity was elevated in HET BR33 iMGs, whereas it was unchanged in HET BR24 iMGs (Extended Data Fig. 4b). Notably, given that both cell lines exhibit reduced lysosomal degradative capacity upon *INPP5D* haploinsufficiency (Fig. 2b), the relatively preserved or increased cathepsin B activity was unexpected. Next, immunostaining for cathepsin B showed a reduction in cathepsin B puncta area but a significant increase in cathepsin B intensity in HET BR24 iMGs, and a trending increase in HET BR33 iMGs (Extended Data Fig. 4c-e). As cathepsin B exhibits a punctate pattern when confined to lysosomes but becomes more diffuse upon release into the cytosol following LMP ^41–43^, the reduction in puncta area despite maintained or elevated enzymatic activity suggests redistribution of cathepsin B from lysosomes to the cytosol in HET iMGs. To directly test this hypothesis, live-cell imaging was performed of iMGs co-stained with LysoTracker and Magic Red. Following development of the Magic Red signal, HET iMGs, similar to iMGs treated with LLoMe, exhibited an increased proportion of cathepsin B activity localized outside LysoTracker+ compartments, consistent with cytosolic redistribution (Figs. 3d-f). We observed a similar redistribution of cathepsin B activity in iMGs treated with the SHIP1-specific inhibitor 3AC^44^ , leading us to hypothesize that SHIP1’s phosphatase activity may be directly involved in its regulation of lysosomal membrane integrity. Consistent with this, 6-hour treatment of iMGs with 3AC dose-dependently reduced LysoTracker intensity without affecting LAMP1+ puncta area or cell viability (Fig. 3g, Extended Data Fig. 4f-g). Together, these data support a model in which SHIP1 phosphatase activity is required to maintain lysosomal membrane integrity.

To investigate whether cytosolic cathepsin B activity underlies the NLRP3 inflammasome activation downstream of reduced SHIP1 activity, iMGs were treated with 3AC, which we previously showed robustly induces the secretion of IL-1β and IL-18 upon activation of the NLRP3 inflammasome^11^. As a positive control, LLoMe treatment activated the NLRP3 inflammasome and increased caspase-3 cleavage, both of which were rescued by pre-treatment with the cathepsin B inhibitor CA-074-Me^45^ (Extended Data Figs. 4h-i).

We next pre-treated WT iMGs with 10 and 200 µM of CA-074-Me for 1 hour prior to the addition of SHIP1 inhibitor (3AC) for 6 hours. CA-074-Me pre-treatment rescued the 3AC-induced IL-1β and IL-18 secretion in a dose-dependent manner (Fig. 3h). Together, these findings support a model in which reduced SHIP1 activity promotes lysosomal membrane permeabilization and release of cathepsin B into the cytosol, thereby driving NLRP3 inflammasome activation (Fig. 3i).

### *INPP5D* haploinsufficiency induces a shift from an immune-responsive state to a DAM-like, phagocytic state in microglia

Endo-lysosomal dysfunction and lipid droplet accumulation have increasingly been linked to disease-associated microglial states in aging and neurodegeneration^35,46,47^. To determine whether the endo-lysosomal defects induced by INPP5D haploinsufficiency were accompanied by alterations in microglial state, we performed CITE-seq on BR33 WT and HET iMGs with a custom panel of microglia-relevant antibodies^48^, which allowed for simultaneous surface epitope and transcriptome profiling at a single cell resolution^49^ (Fig. 4a). After filtering the data to remove suspected doublets and cells with >15% of genes mapping to mitochondrial genes, the remaining dataset had 61,892 cells (29,317 WT iMGs, 32,575 HET iMGs).

**Fig. 4:**
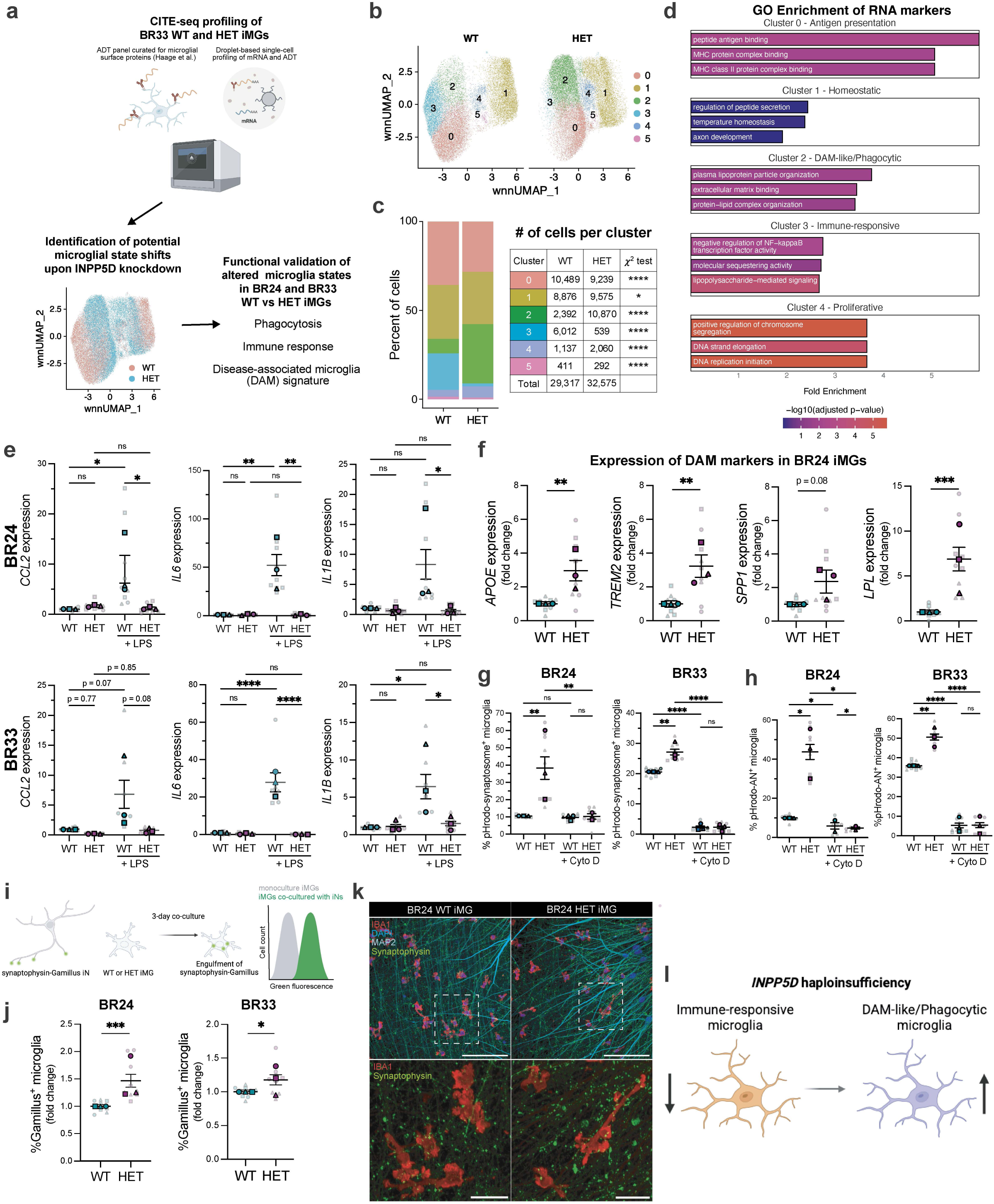
***INPP5D* haploinsufficiency promotes a disease-associated phagocytic program while suppressing immune-response pathways in human microglia** To screen for state shifts with reduction of SHIP1 levels, BR33 WT and HET iMGs, with 2 technical duplicates per condition, were processed on D43 for CITE-seq to simultaneously profile the transcriptome and cell surface proteome, using a custom antibody panel for microglia-relevant antibodies^48^(a-d). Following CITE-seq, key phenotypes were then validated using additional methods in both BR33 and BR24 (e-k). a. Schematic for CITE-seq in BR33 HET and iMGs, overlay of WT and HET iMGs in UMAP generated from integrated RNA and protein (ADT) modalities and functional validation strategy. Created with biorender.com. b. UMAP showing microglia clusters separated by *INPP5D* genotype. c. Relative percentage and number of cells in each cluster in WT vs HET iMGs. Difference in composition between WT and HET iMGs across clusters is calculated using chi-squared test. d. Gene ontology (GO) enrichment results of the positive RNA markers of clusters 1-4. e. mRNA levels of *IL1B*, *IL6* and *CCL2*, measured via RT-qPCR, in BR24 and BR33 WT and HET iMGs treated with LPS (100 ng/mL) or vehicle (water) for 6 hours. f. mRNA levels of DAM markers *APOE*, *TREM2* and *SPP1* and *LPL* measured via RT-qPCR in BR24 WT and HET iMGs. g. Percentage of BR24 and BR33 WT and HET iMGs positive for pHrodo-labeled synaptosomes in the presence or absence of cytochalasin D (cyto D), measured via flow cytometry. h. Percentage of BR24 and BR33 WT and HET iMGs positive for pHrodo-labeled apoptotic neurons (ANs) in the presence or absence of cytochalasin D (cyto D), measured via flow cytometry. i. Experimental strategy for measuring microglial engulfment of synapses attached to live neurons. Created with BioRender.com. j. Percentage of BR24 and BR33 WT and HET iMGs positive for Gamillus after co-culture with synaptophysin-Gamillus iNs, quantified by flow cytometry. k. Representative confocal images of BR24 WT and HET iMGs co-cultured with synaptophysin-Gamillus iNs for 3 days. Scale bars: 100 μm, inset scale: 25 μm. l. Schematic summarizing change in microglia state upon *INPP5D* haploinsufficiency. Unless otherwise stated, data is shown as mean ± SEM, normalized to WT conditions, obtained from n=3 independent differentiations with 3 technical replicates per differentiation. Mixed-effects analysis was performed with genotype and differentiation as fixed factors. The reported p-value corresponds to the main effect of genotype, averaged across differentiations. ****p < 0.0001, ***p < 0.001, **p < 0.01, *p<0.05; ns, not significant. For all scatter plots, different shapes delineate data from separate differentiations, average values per differentiation are shown, with faded dots representing technical replicates (wells) within each differentiation.

We integrated RNA and antibody-derived tag (ADT) data using Seurat’s weighted nearest neighbors (WNN) framework to define cell clusters informed by both modalities. While WT and HET iMGs overlapped to some degree in UMAP space, the cluster compositions were significantly shifted (Figs. 4b-c). Top gene and surface protein markers were identified for each cluster (Extended Data Figs. 5a-c, Supplementary Tables 4-5) and gene ontology (GO) enrichment analysis performed to gain insight into the identities of these clusters (Fig. 4d, Supplementary Table 6). Cluster 0, which is reduced in HET iMGs, shows enrichment (adjusted p-value < 0.05) for MHC-related GO terms, suggesting that this subcluster represents antigen-presenting microglia. Accordingly, the top protein marker for cluster 0 is the antigen-presenting protein CD1c. While cluster 1, which is proportionally similar across WT and HET iMGs, does not have GO terms with adjusted p-value < 0.05, the top terms include regulation of peptide secretion and temperature homeostasis. *P2RY12* and *CX3CR1* are two of the top RNA markers of cluster 1, and SIRP-A is the strongest protein marker, suggesting that homeostatic microglia make up cluster 1. Cluster 2 is significantly expanded in HET iMGs and is enriched in canonical disease-associated microglia (DAM) markers such as *APOE*, *SPP1* and *TREM2*. In line with the previously reported increased phagocytic activity associated with microglia expressing a DAM^47^, the top protein marker of cluster 2 is the Fc gamma receptor CD64. Cluster 3, which is strongly diminished in HET iMGs, is marked by chemokine-encoding genes *CCL2*, *CCL3* and *CCL4*, and shows enrichment for inflammatory signaling-related GO terms. One of the top protein markers of this cluster is CD13, a myeloid receptor that regulates LPS-induced TLR4 signaling^50^. Cluster 4, which differentially expresses cell proliferation-related markers such as *MKI67* and *CDK1*, likely represents proliferative microglia. The minor expansion of this cluster in SHIP1-deficient microglia supports previous reports showing that SHIP1 negatively regulates myeloid cell proliferation^51,52^. Cluster 5 makes up less than 1% of the microglia and does not show significant GO terms nor a positive protein marker, making its identity unclear.

The significant expansion of cluster 2 and reduction of cluster 3 in HET iMGs prompted us to further investigate differential protein expression across WT and HET iMGs within these clusters. Several phagocytic receptors, such as FCER1A, CD32/FCGR2, CD11b and CD33 were significantly upregulated at the cell surface in HET iMGs within cluster 2 (Extended Data Fig. 5d-e, Supplementary Table 7). In contrast, within cluster 3, CD13 (the top marker of cluster 3) was significantly downregulated at the cell surface in HET iMGs, alongside other proteins involved in immune response, such as LAIR1, CCR6 and C5aR (Extended Data Figs. 5d-e, Supplementary Table 7).

The shift in BR33 HET iMGs from an immune-responsive to a DAM-like, phagocytic state observed in the CITE-seq data prompted us to functionally validate these changes in both cell lines. To test whether SHIP1 deficiency impairs NF-κB-mediated response to inflammatory stimuli, as suggested by the reduction of cluster 3, we measured the expression of *IL1B*, *CCL2* and *IL6* at baseline or with LPS treatment via qPCR. HET iMGs did not show a difference in the expression of either gene at baseline relative to WT iMGs. Strikingly, while LPS treatment resulted in increased expression of these genes in WT iMGs as expected, it failed to do so in HET iMGs from both genetic backgrounds (Figs. 4e). The reduced LPS-induced *IL-6* transcription in HET iMGs was paralleled by diminished IL-6 secretion (Extended Data Fig. 5f). Therefore, SHIP1 deficiency impairs NF-κB-dependent response to LPS in microglia.

Next, we validated the increase in expression of the DAM markers *TREM2*, *APOE*, *SPP1* and *LPL* (Fig. 4f). Notably, we observed a significant increase in secreted SPP1 and APOE levels in HET iMGs from both genetic backgrounds (Extended Data Figs. 5g-h), suggesting that the expansion of this cluster has the potential to influence neighboring cells in a non-cell autonomous manner.

To validate effects of *INPP5D* haploinsufficiency on phagocytosis in human microglia, we tested the uptake of various substrates, including synaptic material, apoptotic neurons and Aβ. First, iMGs were treated with pHrodo-labeled synaptosomes for 18 hours and uptake was quantified via flow cytometry (Extended Data Fig. 5i). A significantly higher percentage of HET iMGs took up synaptosomes, which was blocked by treatment with the actin polymerization inhibitor cytochalasin D (Fig. 4g). The percentage of pHrodo-positive HET iMGs remained higher than WT iMGs in the presence bafilomycin A1 (Extended Data Fig. 5j), suggesting that *INPP5D* haploinsufficiency in iMGs increases synaptosome uptake even with induced lysosomal degradation impairment. In addition to synaptosomes, the uptake of pHrodo-labeled apoptotic neurons (ANs) also was tested and found to result in a similar increase in the percentage of pHrodo-positive HET iMGs compared with WT iMG (Fig. 4h).

To investigate whether SHIP1 modulates engulfment of synapses present in living neurons, in addition to isolated synaptic terminals, we co-cultured iMGs with human iPSC-derived neurons (iNs) expressing the fusion protein synaptophysin-Gamillus^53^ and measured Gamillus intensity within iMGs via flow cytometry (Fig. 4i, Extended Data 5k). Gamillus is an acid-tolerant fluorescent protein that does not get quenched in lysosomes and therefore allows for measuring the amount of synaptophysin within the entire endo-lysosomal system of iMGs^54^. We observed a higher percentage of HET iMGs positive for Gamillus as well as higher average Gamillus fluorescence in HET iMGs compared to WT iMGs, indicating that *INPP5D* haploinsufficiency enhances the engulfment of synaptic material from co-cultured neurons (Fig. 4j-k). Moreover, DiI staining revealed a lower number of dendritic spines in iNs co-cultured with HET iMGs (Extended Data Fig. 4l-m). These results suggest that *INPP5D* haploinsufficiency enhances the uptake of multiple lipid-rich substrates, including synaptic material from mature neurons. Collectively, these findings indicate that *INPP5D* haploinsufficiency drives a DAM-like, hyperphagocytic microglial state at the expense of immune responsiveness to stimuli (Fig. 4l).

### Dysregulated TREM2 signaling contributes to both phagocytic and degradative phenotypes in SHIP1-deficient microglia, molecularly linking these two phenotypes

SHIP1’s inhibition of TREM2 signaling has been suggested as one of the mechanisms by which SHIP1 regulates phagocytosis^55,56^. Consistent with prior studies of *Inpp5d* haploinsufficiency in mouse models^57,58^, we observed increased surface expression of TREM2 in HET iMGs (Fig. 5a), supporting previous work showing enhanced TREM2 signaling upon reduced SHIP1 activity.

**Fig. 5:**
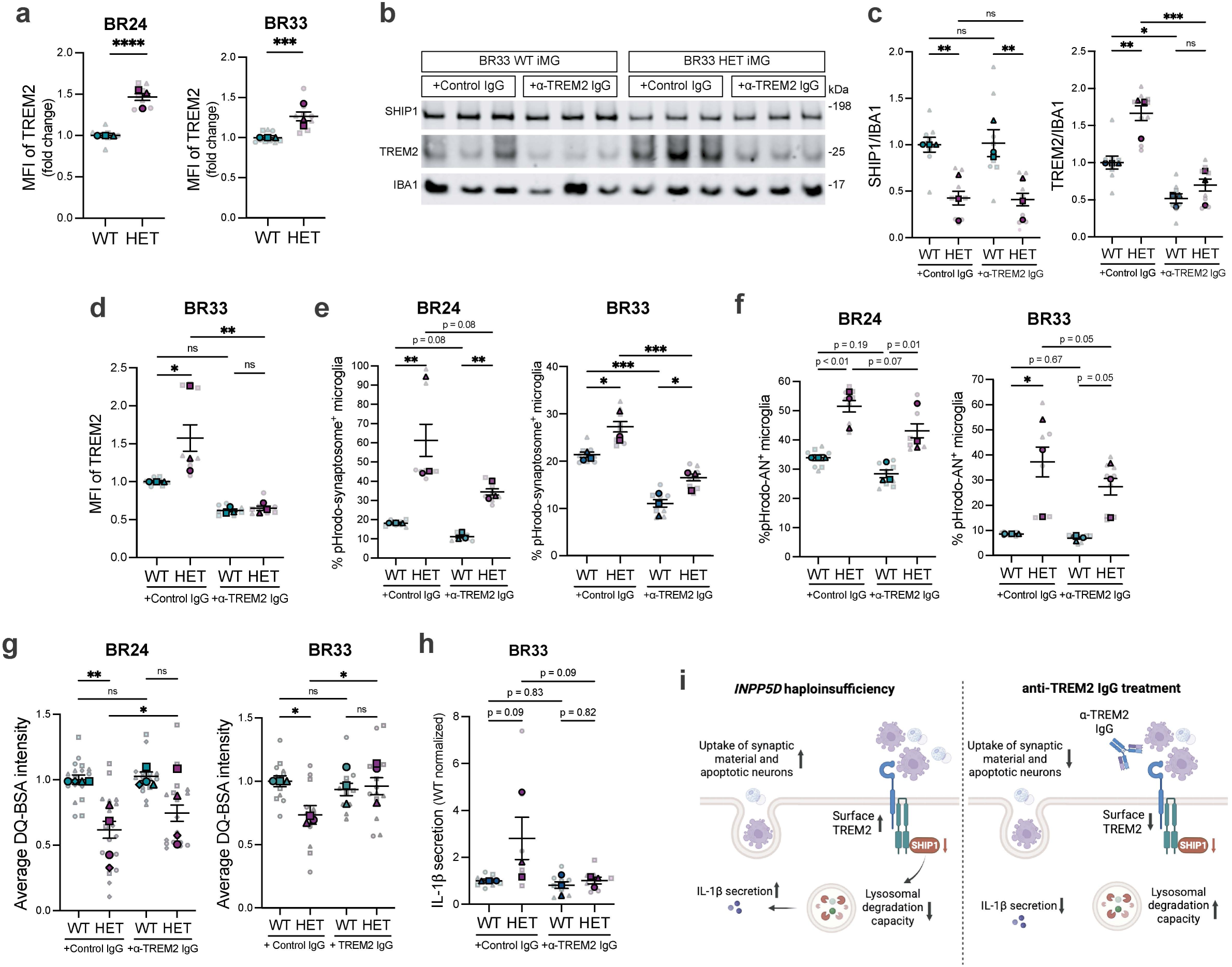
Anti-TREM2 antibody treatment suppresses excessive phagocytosis and improves lysosomal function in SHIP1-deficient microglia. a. MFI of surface TREM2 in non-permeabilized live BR24 and BR33 WT and HET iMGs, measured by flow cytometry. b. Representative Western blot of BR33 WT and HET iMGs treated with control or anti-TREM2 IgG. c. Western blot quantification of SHIP1 and TREM2, normalized by IBA1, in BR33 WT and HET iMGs. Two-way ANOVA was performed using averages from each differentiation. d. MFI of surface TREM2 in non-permeabilized live BR33 WT and HET iMGs, treated with control or TREM2 IgG overnight, measured by flow cytometry. Two-way ANOVA was performed using averages from each differentiation. e. Percentage of BR24 and BR33 WT and HET iMGs positive for pHrodo-synaptosomes and (f) pHrodo-labeled ANs after treatment with control or anti-TREM2 IgG overnight, measured via flow cytometry. Two-way ANOVA was performed using averages from each differentiation. g. Average DQ-BSA intensity, measured via IncuCyte, in BR24 and BR33 WT and HET iMGs treated with control or anti-TREM2 IgG for 3 days. Two-way ANOVA was performed using averages from each differentiation. h. IL-1β secretion in BR33 WT and HET iMGs treated with control or anti-TREM2 IgG for 3 days. i. Proposed model of effect of anti-TREM2 antibody treatment in the context of *INPP5D* haploinsufficiency. Created with BioRender.com. Unless otherwise stated, data is shown as mean ± SEM, normalized to WT conditions, obtained from n=3 independent differentiations with 3 technical replicates per differentiation. Mixed-effects analysis was performed with genotype and differentiation as fixed factors. The reported p-value corresponds to the main effect of genotype, averaged across differentiations. ****p < 0.0001, ***p < 0.001, **p < 0.01, *p<0.05; ns, not significant. For all scatter plots, different shapes delineate data from separate differentiations, average values per differentiation are shown, with faded dots representing technical replicates (wells) within each differentiation.

To directly assess the contribution of TREM2 to the phenotypes associated with SHIP1 deficiency in human microglia, iMGs were treated with an anti-TREM2 antibody (R&D Systems, AF18281). This treatment reduced both total and surface TREM2 levels in HET iMGs compared to WT iMGs, without affecting SHIP1 levels (Figs. 5b–d), confirming effective normalization of TREM2 levels independent of SHIP1.

Functionally, anti-TREM2 antibody treatment suppressed the elevated uptake of synaptosomes and ANs in HET iMGs (Fig. 5e), indicating that increased TREM2 signaling contributes to the

hyperphagocytic phenotype. Notably, anti-TREM2 treatment also ameliorated the lysosomal degradation deficit and reduced the elevated IL-1β secretion observed in HET iMGs (Figs. 5f-g), suggesting that rescue of TREM2 levels may also rescue lysosomal deficits and inflammation induced by reduction of SHIP1 in human microglia. Together, these findings implicate dysregulated TREM2 signaling in driving excessive phagocytic uptake, impaired lysosomal processing, and increased IL-1β secretion in SHIP1-deficient microglia (Fig. 5h).

### *INPP5D* haploinsufficiency does not affect Aβ uptake but leads to Aβ accumulation in microglia

Previous studies in mouse models and immortalized microglia cell lines suggested a role for SHIP1 in regulating Aβ phagocytosis^59–61^. We tested the effect of microglial *INPP5D* haploinsufficiency on Aβ uptake via flow cytometry of iMGs treated with 0.1 μM FITC-labeled fibrillar Aβ_42_ (FITC-fAβ) for 18 hours. *INPP5D* haploinsufficiency did not affect the percentage of iMGs positive for Aβ or the fluorescence intensity of Aβ within iMGs (Figs. 6a-b, Extended Data Fig. 6a).

**Fig. 6:**
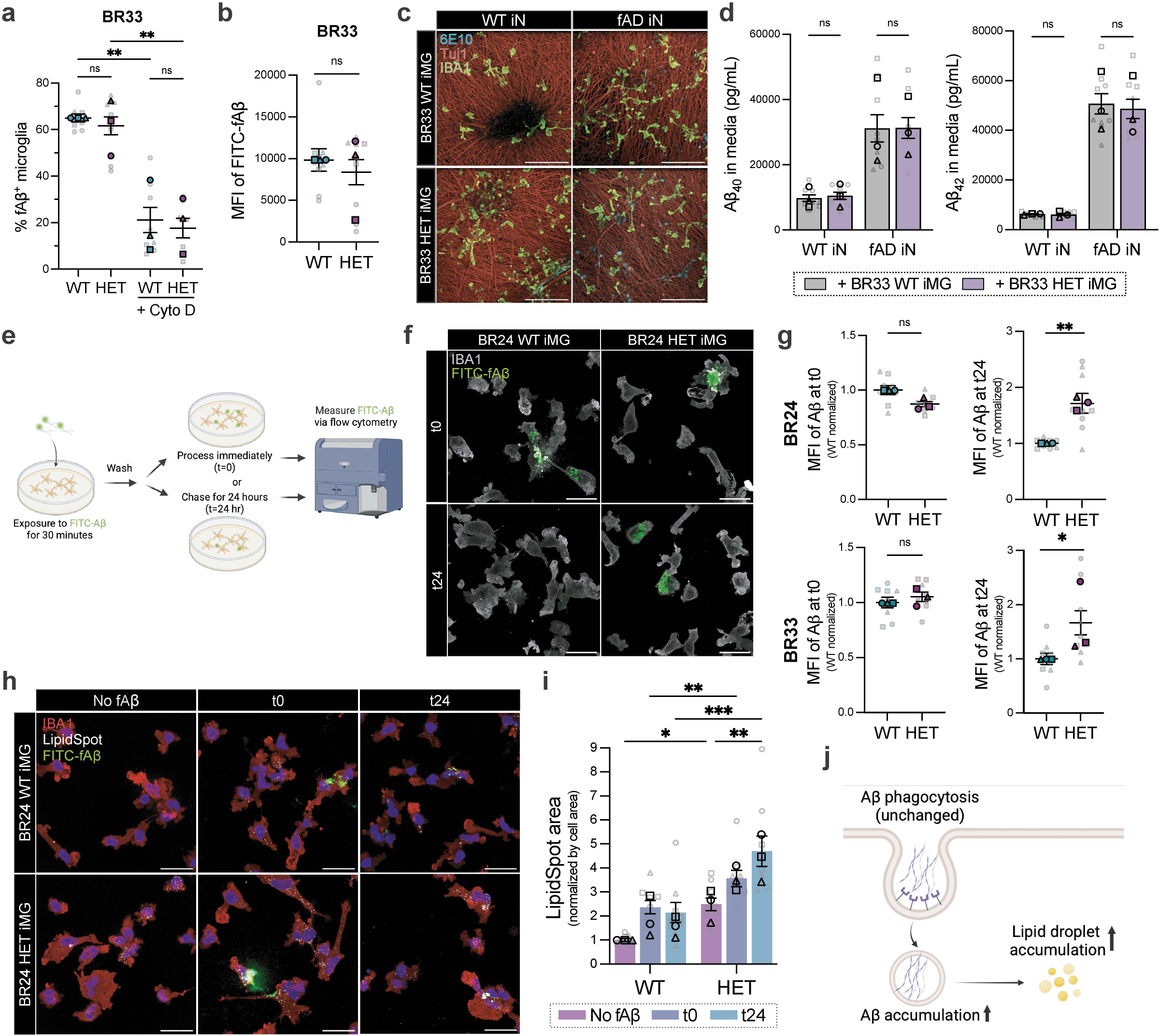
***INPP5D* haploinsufficiency does not affect Aβ uptake but leads to Aβ accumulation in microglia** a. Percentage of BR33 WT and HET iMGs positive for FITC-labeled fibrillar Aβ (fAβ) in the presence or absence of cyto D, measured via flow cytometry. Two-way ANOVA was performed using averages from each differentiation b. MFI of FITC-fAβ in non-permeabilized, live BR33 iMGs. c. Representative confocal images of iMG-iN co-cultures immunostained for 6E10, IBA1 and Tuj1. Scale bars: 100 μm. d. Aβ_40_ and Aβ_42_ measured in the conditioned media of iN-iMG co-cultures via ELISA. Quantification shown for n=3 differentiations with 3 technical replicates per differentiation. See Two-way ANOVA was performed using averages from each differentiation. e. Experimental scheme for measuring the accumulation of FITC-fAβ in iMGs. Created with BioRender.com. f. Representative confocal images of BR24 iMGs at after 30 minutes of FITC-fAβ uptake (t0) or 24 hours post-chase after FITC-fAβ treatment (t24). Scale bars: 25 μm. g. MFI of FITC-fAβ at t0 (left) and t24 hour in BR24 and BR33 iMGs. See Extended Data Fig. 6c for IncuCyte quantification of FITC-fAβ intensity within microglia within the same wells over chase time. h. Representative confocal images of BR24 WT and HET iMGs without Aβ treatment, at t0 and at t24 counterstained with LipidSpot. Scale bars: 25 μm. i. Quantification of LipidSpot puncta area normalized by IBA1^+^ area. Two-way ANOVA with Tukey’s multiple comparisons test was performed using averages from each differentiation. j. Schematic summarizing findings in figure 6. Created with BioRender.com Unless otherwise stated, data is shown as mean ± SEM, normalized to WT conditions, obtained from n=3 independent differentiations with 3 technical replicates per differentiation. Mixed-effects analysis was performed with genotype and differentiation as fixed factors. The reported p-value corresponds to the main effect of genotype, averaged across differentiations. ****p < 0.0001, ***p < 0.001, **p < 0.01, *p<0.05; ns, not significant. For all scatter plots, different shapes delineate data from separate differentiations, average values per differentiation are shown, with faded dots representing technical replicates (wells) within each differentiation.

To investigate whether SHIP1 regulates the uptake of cell-derived Aβ, we co-cultured iMGs with iNs carrying homozygous familial Alzheimer’s disease (fAD) mutations in Amyloid Precursor Protein and Presenilin 1 (APP^Swe/Swe^; PSEN1^M146V/M146V^) and a WT isogenic control line (WT )^62,63^. fAD iNs secreted elevated Aβ_42_ and an increased ratio of Aβ_42_/Aβ_40_, compared to paired isogenic WT iNs, as expected (Extended Data Fig. 6b). Aβ_42_ and Aβ_40_ remaining in the media of iMG-iN co-cultures was measured via ELISA, and clearance of extracellular Aβ was not measurably affected by the presence of HET compared to WT iMGs (Figs. 6c-d). Taken together, these data do not suggest a role for SHIP1 in regulating Aβ uptake in this context.

Next, we interrogated whether *INPP5D* haploinsufficiency leads to an accumulation of phagocytosed Aβ within microglia. WT and HET iMGs were treated with FITC-fAβ for 30 minutes, after which excess Aβ from the media was removed and the remaining FITC fluorescence in iMGs was quantified after 24 hours via flow cytometry (Figs. 6e-f). HET iMGs from both genetic backgrounds displayed higher FITC fluorescence 24 hours after washout compared to WT iMGs, suggesting an intracellular accumulation of Aβ (Fig. 6g, Extended Data Fig. 6c). Moreover, Aβ accumulation over 24 hours exacerbated the lipid droplet accumulation in HET, but not WT, iMGs (Figs. 6h-i). Together, these data suggest that while *INPP5D* haploinsufficiency does not affect Aβ uptake, it leads to the accumulation of phagocytosed Aβ within microglia, likely due to impaired lysosomal degradation. Impaired Aβ degradation by HET microglia is accompanied by excess lipid droplet accumulation, which is already elevated at baseline, thereby potentially exacerbating neuroinflammation (Fig. 6j).

Across two independent genetic backgrounds, *INPP5D* haploinsufficiency produced highly concordant endo-lysosomal, inflammatory, and cargo-selective phagocytic phenotypes in human iMGs (Extended Data Fig. 7). We therefore next investigated whether similar cellular programs were associated with endogenous variation in SHIP1 abundance across genetically diverse human microglia.

### The protective allele of the AD-associated *INPP5D* SNP rs10933431 is associated with enhanced endo-lysosomal trafficking pathways and increased abundance of catalytically active SHIP1 species

To determine whether the cellular changes identified in SHIP1-deficient microglia are associated with endogenous variation in SHIP1 abundance across a genetically diverse human population, we generated iMGs from 75 donors from the ROSMAP cohort^17,64^ and performed proteomic profiling using data-independent acquisition mass spectrometry (Figs. 7a-b, Supplementary table 8). We first examined the correlation of SHIP1 levels proteome-wide to identify pathways associated with endogenous variation in SHIP1 abundance (Fig. 7c, Supplementary table 9). Gene set enrichment analysis (GSEA) of SHIP1-associated proteins revealed positive enrichment of lysosome and phagocytosis pathways (Fig. 7d, Supplementary table 10), and SHIP1 abundance positively correlated with GSVA scores for proteins in the KEGG “Lysosome” pathway across donor-derived iMGs (Fig. 7e), further supporting an association between SHIP1 levels and endo-lysosomal activity in human microglia. Selected GO biological process terms from the GSEA demonstrated enrichment of pathways related to phagocytosis, endocytosis, protein localization to endosomes, endosomal transport, and endosome-to-lysosome trafficking among SHIP1-correlated proteins (Fig. 7f, Supplementary table 11). Together, these findings demonstrate that endo-lysosomal and phagocytic pathways associated with SHIP1 deficiency are also linked to endogenous variation in SHIP1 abundance across genetically diverse human microglia.

**Fig. 7:**
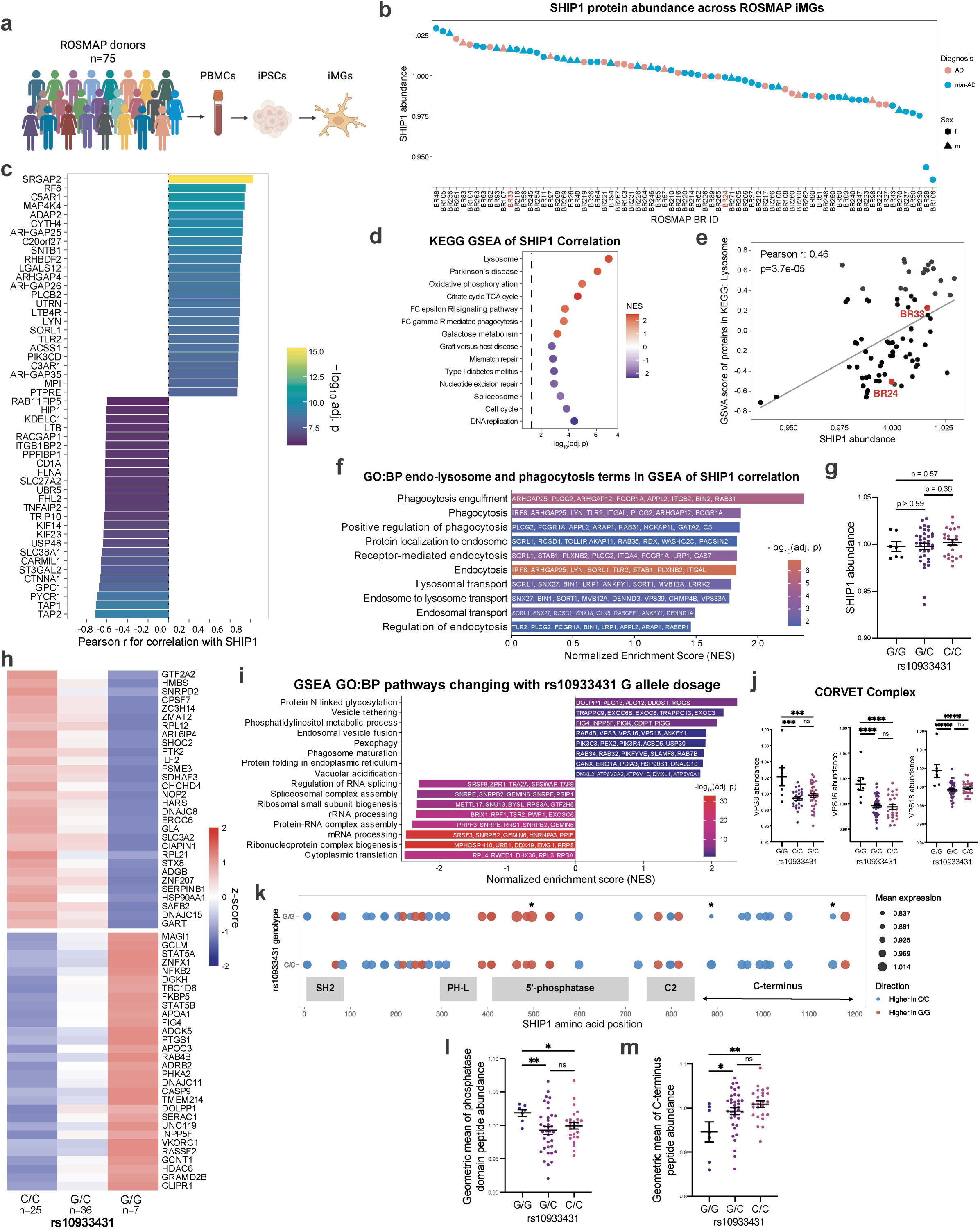
SHIP1 abundance and the protective *INPP5D* rs10933431 allele are associated with enhanced endolysosomal trafficking programs in human microglia. a. Experimental schematic of ROSMAP donor-derived iMG generation. Created with BioRender.com. See Supplementary Table 8 for meta data of the donor lines. b. SHIP1 protein abundance across ROSMAP donor-derived iMGs colored by AD diagnosis and sex. c. Top 25 proteins positively or negatively correlated with SHIP1 abundance across donor-derived iMGs, ranked by Pearson correlation. See Supplementary Table 9 for correlation of all proteins with SHIP1. d. KEGG gene set enrichment analysis (GSEA) of proteins correlated with SHIP1 abundance. See Supplementary Table 10 for all KEGG GSEA results. e. Correlation between SHIP1 abundance and gene set variation analysis (GSVA) scores for proteins in the KEGG Lysosome pathway across donor-derived iMGs. f. Selected GO biological process pathways enriched in GSEA of SHIP1-correlated proteins related to endo-lysosomes and phagocytosis. See Supplementary Table 11 for all GO GSEA results. g. Total SHIP1 abundance across rs10933431 genotypes. One-way ANOVA with Fisher’s LSD post hoc test. h. Heatmap of differentially abundant human proteins with p-value<0.05 across rs10933431 genotypes ordered by G allele dosage. See Supplementary Table 12 for all differentially abundant proteins. i. Selected GO biological process pathways enriched in GSEA of proteins associated with rs10933431 G allele dosage. See Supplementary Table 13 for all GO GSEA results. j. Abundance of CORVET complex proteins VPS8, VPS16, and VPS18 across rs10933431 genotypes. One-way ANOVA with Fisher’s LSD post hoc test. k. Mapping of detected SHIP1 proteomic peptides across the SHIP1 protein structure by rs10933431 genotype. Dot size indicates mean peptide abundance and color indicates relative enrichment between G/G and C/C genotypes. Asterisks indicate peptides significantly differing between genotypes determined by two-sided Wilcoxon rank-sum test. l. Geometric mean abundance of peptides mapping to the SHIP1 5′-phosphatase domain across rs10933431 genotypes. One-way ANOVA with Fisher’s LSD post hoc test. m. Geometric mean abundance of peptides mapping to the SHIP1 C-terminus across rs10933431 genotypes. One-way ANOVA with Fisher’s LSD post hoc test. ****p < 0.0001, ***p < 0.001, **p < 0.01, *p<0.05; ns, not significant. Each data point represents a genetic background.

Given the association between SHIP1 abundance and endo-lysosomal pathway activity across donor-derived iMGs, we next investigated whether the AD-associated *INPP5D* SNP rs10933431 associates with altered SHIP1 abundance or endolysosomal programs in human microglia. The minor allele G of rs10933431 was identified as protective against AD risk with an odds ratio of 0.93 in a large GWAS meta-analysis^8^. The rs10933431 genotype distribution in our dataset consisted of 25 C/C, 36 G/C, and 7 G/G individuals (n = 68), corresponding to a protective G-allele frequency of 36.8%. This frequency is higher than that reported in European ancestry reference populations (23.4%)^8^, improving our ability to identify allele-dosage-dependent changes in the microglial proteome.

Consistent with previous reports ^12,65^, we did not observe a difference in SHIP1 abundance across rs10933431 genotypes (Fig. 7g). We therefore investigated whether rs10933431 genotype associates with broader proteomic changes in human microglia by performing an allele-dosage analysis across donor-derived iMGs, modeling rs10933431 genotype as 0, 1, or 2 copies of the protective G allele while adjusting for sex and ancestry (Fig. 7h, Supplementary table 12). Proteins increasing with rs10933431-G allele dosage were enriched in pathways such as “phosphatidylinositol metabolic process”, “vesicle tethering”, “endosomal vesicle fusion” and “phagosome maturation”, whereas proteins decreasing with G allele dosage were associated with RNA processing (Fig. 7i, Supplementary table 13). Notably, multiple components of the CORVET complex, an early endosomal tethering complex involved in Rab5-dependent endosomal fusion and maturation including VPS8, VPS16 and VPS18, were increased in protective G/G carriers compared to non-protective C allele carriers (Fig. 7j). Together, these findings support an association between the protective allele of rs10933431 and endo-lysosomal trafficking programs in human microglia.

Given the enrichment of phosphatidylinositol metabolic pathways with increasing rs10933431-G allele dosage, we next mapped the SHIP1 peptides detected by proteomics onto the SHIP1 protein structure to determine whether the protective allele is associated with altered abundance of specific SHIP1 domains (Fig. 7k). While total SHIP1 protein abundance did not differ detectably across rs10933431 genotypes (Fig. 7g), peptides mapping to the SHIP1 5′-phosphatase domain were increased in protective G/G genotypes compared to G/C and C/C genotypes (Fig. 7l). In contrast, peptides mapping to the SHIP1 C-terminus were reduced in protective G/G carriers relative to non-protective allele carriers (Fig. 7m). These findings are consistent with previous reports demonstrating reduced SHIP1 phosphatase-domain abundance and increased C-terminal abundance in AD brain tissue^11,12^. Together, these findings raise the possibility that increased abundance of catalytically active SHIP1 species in protective rs10933431 carriers contributes to enhanced endo-lysosomal trafficking programs in human microglia and a beneficial effect on AD risk.

### High Aβ in the brain amplifies dysfunctional phenotypes in SHIP1-deficient human microglia *in vivo*

To determine whether the Alzheimer’s pathology in the brain exacerbates the dysfunctional phenotypes observed in SHIP1-deficient human microglia *in vivo*, WT or *INPP5D* haploinsufficient iPSC-derived hematopoietic microglial progenitors (hematopoietic progenitor cells; HPCs) were xenotransplanted into 2 month-old adult WT-hFIRE or 5xFAD-hFIRE (5x-hFIRE) mice^64,66^. Importantly, hFIRE and 5x-hFIRE mice are Rag2- and Il2rγ-deficient, genetically express humanized CSF1, and develop without microglia allowing for CNS-wide human microglial engraftment. Analysis of brain sections collected at 6 months of age found WT and HET xenografted iMGs (xMG) engrafted efficiently in both WT-hFIRE and 5x-hFIRE brains (Fig. 8a-b, Extended Data Figs. 8a-c). Despite previous reports suggesting that SHIP1 regulates amyloid plaque clearance in mouse models, *INPP5D* haploinsufficiency in xenotransplanted iMGs (xMGs) did not detectably alter amyloid plaque burden in 5x-hFIRE brains (Extended Data Figs. 8d-e), consistent with our *in vitro* findings demonstrating unaltered Aβ uptake by HET iMGs (Figs. 6a-d).

**Fig. 8:**
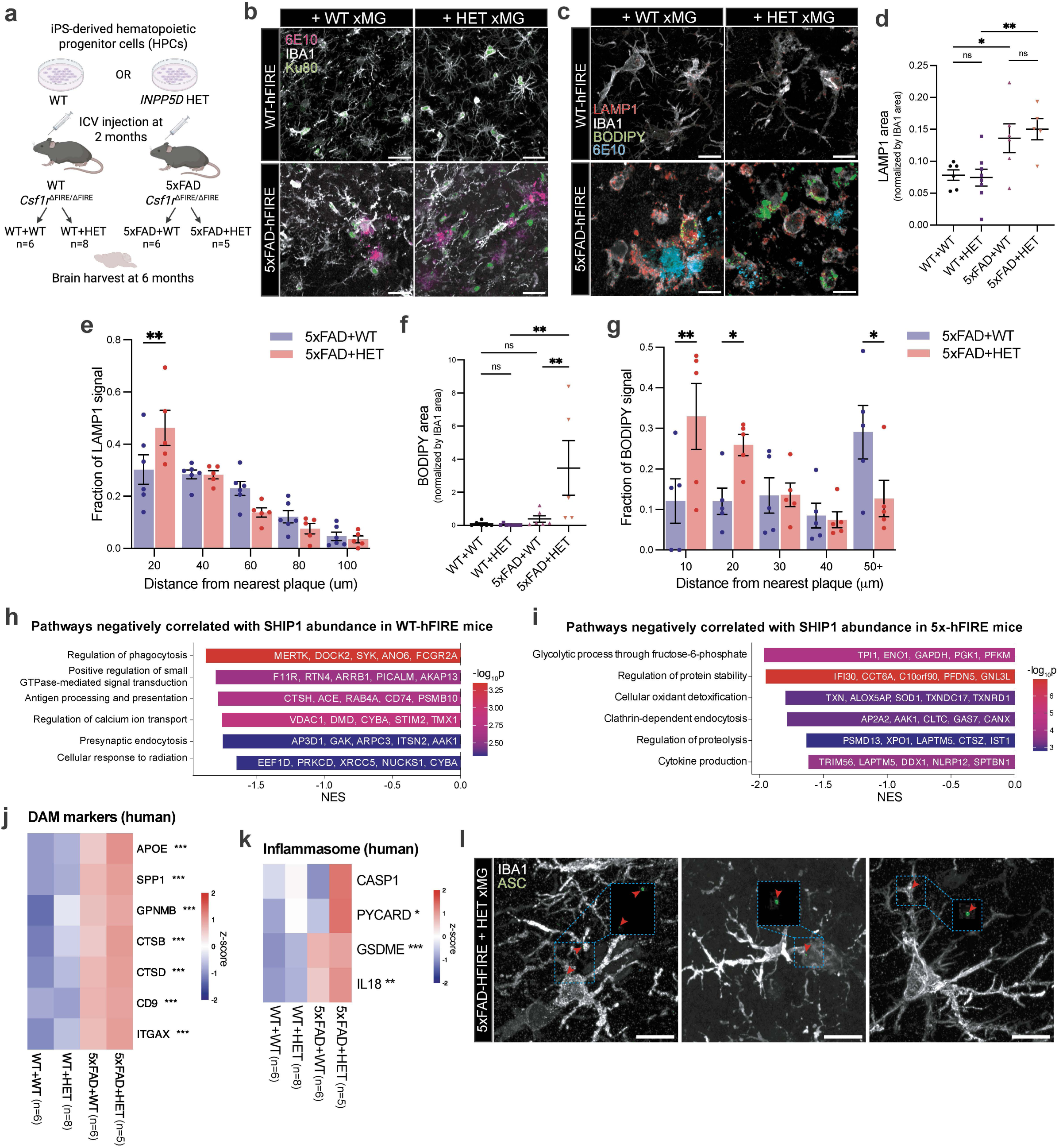
5x-hFIRE brain environment amplifies lysosomal, lipid droplet, and inflammatory phenotypes in SHIP1-deficient human microglia. a. Experimental schematic of xenotransplantation of BR33 WT or *INPP5D* haploinsufficient (HET) iPSC-derived HPCs into 2-month-old adult WT-hFIRE or 5x-hFIRE mice followed by analysis of xenografted human microglia at 6 months. b. Representative confocal images of engrafted human xMGs (Ku80+, IBA1+) surrounding 6E10-positive plaques in WT-hFIRE and 5xFAD-hFIRE brains. Scale bars: 25 μm. c. Representative confocal images of LAMP1-positive lysosomal structures and BODIPY-positive lipid droplets in engrafted human xMGs surrounding 6E10+ plaques. Scale bars: 25 μm. d. Quantification of LAMP1-positive area normalized to IBA1 area across experimental groups. e. Spatial quantification of LAMP1 signal as a function of distance from the nearest amyloid plaque in xMGs engrafted into 5x-hFIRE mice. f. Quantification of BODIPY-positive area normalized to IBA1 area across experimental groups. g. Spatial quantification of BODIPY signal as a function of distance from the nearest 6E10+ plaque in xMGs engrafted into 5x-hFIRE mice. h. Selected GO biological process pathways negatively correlated with SHIP1 abundance in human proteins derived from xenografted WT-hFIRE brains. Asterisks indicate proteins significantly differing across groups by one-way ANOVA. See Supplementary Table 16 all GO GSEA results. i. Selected GO biological process pathways negatively correlated with SHIP1 abundance in human proteins derived from xenografted 5x-hFIRE brains. See Supplementary Table 16 all GO GSEA results. j. Heatmap of DAM-associated human proteins across experimental groups. Asterisks indicate proteins significantly differing across groups by one-way ANOVA. k. Heatmap of inflammasome-associated human proteins across experimental groups. Asterisks indicate proteins significantly differing across groups by one-way ANOVA. l. Representative images of ASC-positive puncta in HET human xMGs engrafted into 5x-hFIRE brains. Arrowheads indicate ASC-positive puncta. Scale bars: 10 μm. ****p < 0.0001, ***p < 0.001, **p < 0.01, *p<0.05; ns, not significant. Each data point represents a mouse.

Consistent with previous reports demonstrating increased lysosomal signatures in plaque-associated microglia in 5xFAD mice ^46,47^, xMGs engrafted into 5x-hFIRE brains displayed increased accumulation of LAMP1-positive puncta compared to xMGs engrafted into WT-hFIRE brains (Figs. 8c-d). Spatial analysis revealed that LAMP1 signal was enriched in xMGs proximal to amyloid plaques, with HET xMGs displaying significantly greater plaque-associated LAMP1 intensity within 20 µm of 6E10+ deposits compared to WT xMGs (Fig. 8e). HET xMGs in 5x-hFIRE brains also exhibited markedly increased accumulation of BODIPY-positive lipid droplets concentrated near plaques compared to WT xMGs (Figs. 8f-g), consistent with the lipid droplet accumulation observed following Aβ exposure in HET iMGs *in vitro* (Figs. 6h-i) and previous reports describing lipid-droplet-accumulating dysfunctional microglia in neurodegenerative conditions^35,36^. Together, these findings suggest that high amyloid burden amplifies lysosomal abnormalities associated with SHIP1 deficiency, resulting in the accumulation of lysosome- and lipid-rich material within plaque-associated microglia.

To determine how the brain environment alters SHIP1-associated cellular programs in engrafted human microglia, we performed proteomic profiling on xenografted WT-hFIRE and 5x-hFIRE brains and analyzed human-specific proteins derived from iMGs. Proteins whose abundance significantly correlates with SHIP1 in WT or 5xFAD brain environments are shown (Extended Data Fig. 8f), along with disease-context-dependent SHIP1-associated pathways (Figs. 8h-i, Extended Data Fig. 8g-h, Supplementary table 14-15). In both WT-hFIRE and 5x-hFIRE brains, lower SHIP1 abundance was associated with endocytosis-related pathways, consistent with the endolysosomal defects observed in SHIP1-deficient iMGs. In WT-hFIRE brains, reduced SHIP1 abundance was additionally associated with pathways related to phagocytosis and antigen presentation (Fig. 8h, Supplementary table 16), whereas in 5x-hFIRE brains, lower SHIP1 abundance was associated with pathways related to cytokine production, oxidative detoxification, proteolysis, and glycolytic metabolism (Fig. 8i, Supplementary table 17), suggesting that the 5xFAD brain environment amplifies inflammatory and metabolic dysfunction associated with reduced SHIP1 abundance. To further define proteomic changes associated with *INPP5D* haploinsufficiency *in vivo*, we next performed differential expression analysis comparing human proteins from WT and HET xMGs within WT and 5xFAD brain environments (Extended Data Fig. 9a-b, Supplementary table 18-19). Consistent with the correlation analyses, HET xMGs in WT-hFIRE brains exhibited enrichment of pathways related to antigen presentation, phagocytosis, and immune effector processes, alongside downregulation of phosphatidylinositol metabolic pathways (Extended Data Fig. 9c-d, Supplementary table 20). In 5x-hFIRE brains, HET xMGs demonstrated enrichment of pathways related to oxidative detoxification, proteolysis, protein stabilization, and inflammatory responses (Extended Data Fig. 9e-f, Supplementary table 21), further supporting that the 5xFAD brain environment amplifies dysfunctional inflammatory and metabolic phenotypes associated with SHIP1 deficiency.

We next asked whether the SHIP1-associated proteomic changes in 5x-hFIRE brains were accompanied by disease-associated and inflammasome-related microglial phenotypes. Human proteins from 5x-hFIRE-engrafted brains showed increased abundance of DAM-associated markers, including APOE, SPP1, GPNMB, CTSB, CTSD, CD9, and ITGAX, with the highest levels observed in 5x-hFIRE brains engrafted with HET xMGs (Fig. 8j). Similarly, inflammasome-associated proteins, including CASP1, PYCARD and IL18, were increased in the 5x-hFIRE brains engrafted with HET xMGs (Fig. 8k). Consistent with this proteomic signature, we detected ASC-positive puncta in HET xMGs engrafted into 5x-hFIRE brains (Fig. 8l), suggesting that the plaque-rich 5xFAD brain environment amplifies DAM-like and inflammasome-associated phenotypes in SHIP1-deficient human microglia.

## DISCUSSION

Over the past decade, SHIP1 has drawn increasing attention because its encoding gene, *INPP5D*, has emerged as a genetic risk factor for Alzheimer’s disease^7^. Recently, our group reported that while the mRNA expression of *INPP5D* is increased in brain tissue of individuals with AD, levels of full-length SHIP1 protein that include its phosphatase domain are reduced^67^. In parallel, another group reported an increase in transcripts encoding truncated forms of SHIP1 that lack the phosphatase domain in AD brain tissue^12^. Here, we extend these findings by showing that the risk allele of AD-associated *INPP5D* SNP rs10933431 is associated with decreased abundance of SHIP1 phosphatase domain peptides and increased abundance of C-terminal SHIP1 peptides in human microglia, similar to the pattern observed in AD brain tissue. These findings emphasize the critical need for elucidating the effects of SHIP1 deficiency on microglia cell function and contributions to AD pathogenesis.

SHIP1 is best known as a negative regulator of PI3K/AKT signaling in hematopoietic cells, where it limits inflammatory signaling and immune activation through interactions with adaptor proteins such as SHC1, DOK2, and GRB2^19,20^. Here, we confirmed interactions between SHIP1 and canonical signaling adaptors in human microglia while additionally identifying binding partners associated with the endo-lysosomal machinery. Among these, the CapZ complex is notable, as it has been shown to regulate endosomal trafficking by controlling F-actin density around early endosomes and recruiting Rab5 effectors^24^. Consistent with these observations, SHIP1 deficiency in iMGs altered Rab5+ and Rab7+ endosomal compartments, increased Galectin-3+ puncta number, and reduced lysosomal degradation of internalized cargo, suggesting that SHIP1 contributes to degradative endo-lysosomal trafficking in human microglia. Importantly, proteins associated with vesicle tethering, endosomal vesicle fusion, vacuolar acidification, and phagosome maturation positively correlated with SHIP1 abundance and with the protective rs10933431 allele in genetically diverse human microglia. Together, these findings support a role for SHIP1 in maintaining endo-lysosomal trafficking in human microglia. Future studies are warranted to determine if SHIP1’s interaction with CapZ plays a role in SHIP1’s regulation of endo-lysosomal trafficking.

We additionally identified SHIP2 as an endogenous binding partner of SHIP1. SHIP1 and SHIP2 share conserved SH2 and 5′-phosphatase domains, raising the possibility that they cooperatively regulate phosphoinositide signaling at endolysosomal membranes. Lysosomal membranes contain phosphoinositides that regulate lysosome function and dynamics. Notably, SHIP1 and SHIP2’s enzymatic product, PI(3,4)P_2_, is found on lysosomal membranes, where it controls lysosome function^68^, while their enzymatic substrate, PI(3,4,5)P_3_, can induce lysosome de-acidification by activating the lysosomal potassium channel TMEM175^69^. We found that acute inhibition of SHIP1’s phosphatase activity robustly destabilized lysosomes, further supporting the idea that SHIP1 maintains lysosomal homeostasis via its phosphatase activity.

SHIP1-deficient microglia from both genetic backgrounds exhibited lysosomal stress and membrane destabilization. In *INPP5D* haploinsufficient iMGs, cathepsin B levels and enzymatic activity appeared elevated, whereas cathepsin B+ puncta, which likely represent lysosomally localized cathepsin B, were reduced. These findings suggest that defective degradative trafficking may impair cathepsin delivery to lysosomes and/or promote leakage of active cathepsin B into the cytosol following lysosomal membrane permeabilization^41,70^. In line with this model, both chronic SHIP1 deficiency and acute SHIP1 inhibition increased galectin-3 puncta formation and cathepsin B activity outside lysosomal compartments, indicating impaired lysosomal membrane integrity. Based on the previous literature linking lysosomal membrane permeabilization and cathepsin release to NLRP3 inflammasome activation ^39,40,43^, we reasoned that cathepsin B leaking from destabilized lysosomes may mediate inflammasome activation downstream of SHIP1 deficiency ^11^. Indeed, inhibition of cathepsin B activity with CA-074-Me suppressed IL-1β and IL-18 release following lysosomal damage or SHIP1 inhibition, supporting a role for cathepsin B in mediating NLRP3 inflammasome activation downstream of impaired endolysosomal trafficking in SHIP1-deficient microglia.

Efficient degradation of phagocytic cargo sustains phagocytic uptake in macrophages^71,72^. Accordingly, impaired lysosomal degradation in SHIP1-deficient microglia would be expected to limit further phagocytosis to ensure the degradation of material already internalized. However, surprisingly, SHIP1 deficiency has been associated with *increased* phagocytosis in multiple systems^73,74^. Most recently, SHIP1 deficiency has been shown to increase developmental synapse pruning in mice^75^. In line with these findings, we found that SHIP1 deficiency enhances the uptake of synaptic material and apoptotic neurons, both lipid-rich substrates, in human microglia. We propose that this increased phagocytosis *despite impaired degradative trafficking* is due to altered regulation of phagocytic receptors and induction of a disease-associated phagocytic microglial state following *INPP5D* haploinsufficiency. One mechanism contributing to this phenotype involves increased TREM2 signaling, consistent with previous studies linking SHIP1 deficiency to enhanced TREM2-dependent microglial activation^56,58^. Antibody-mediated modulation of TREM2 partially rescued excessive phagocytosis and improved lysosomal degradation in SHIP1-deficient iMGs (Figs. 5e-g), suggesting that dysregulated TREM2 signaling contributes to both the hyper-phagocytic and degradative phenotypes induced by SHIP1 deficiency, mechanistically linking these two SHIP1 HET phenotypes. These findings raise the possibility that therapeutic modulation of TREM2 signaling may mitigate endo-lysosomal dysfunction in the context of SHIP1 deficiency^77,78^.

SHIP1’s precise role in Aβ clearance has remained difficult to define. Previous mouse studies variably reported reduced, increased, or unchanged plaque burden following *Inpp5d* perturbation^57,58,60,76^, likely reflecting differences in disease stage, mouse model, experimental design, and potentially species-specific effects of SHIP1 deficiency. In this study, we did not find evidence supporting a role for SHIP1 in regulating either synthetic or neuron-derived Aβ uptake in human iMGs. Consistent with these findings *in vitro*, xenografted SHIP1-deficient human microglia did not alter amyloid plaque burden or soluble Aβ levels in 5xhFIRE brains *in vivo*. However, while SHIP1-deficient microglia in vitro also didn’t show a defect in uptake of Aβ, they did display accumulated intracellular Aβ following uptake, consistent with impaired degradative processing rather than altered uptake itself. Moreover, SHIP1-deficient human microglia in 5x-hFIRE brains exhibited increased plaque-associated LAMP1 accumulation and exaggerated lipid droplet accumulation, supporting the hypothesis that amyloid pathology amplifies impaired degradative trafficking in the context of SHIP1 deficiency. This distinction between uptake and intracellular degradation may reconcile prior conflicting findings regarding plaque clearance. Although TREM2 facilitates uptake of lipidated Aβ species^77,78^, altered expression of additional Aβ-binding receptors may compensate for increased TREM2 signaling following SHIP1 deficiency. Moreover, Aβ can associate with APOE and clusterin (CLU) in the brain^78^, potentially altering the receptor pathways involved in its clearance. Together, these findings suggest that SHIP1 primarily regulates intracellular degradative handling of phagocytosed Aβ rather than its uptake.

Endo-lysosomal dysfunction has emerged as a potential key driver of late-onset AD etiology, and recent evidence suggests that Aβ may accumulate intracellularly prior to extracellular plaque deposition^79–81^. In this study, we identify SHIP1 as a central regulator of convergent microglial mechanisms implicated in AD pathogenesis. SHIP1 deficiency recapitulated a number of microglial phenotypes closely associated with AD pathology (see Extended Data Fig. 7 for summarized results). These include aberrant synapse engulfment, intracellular accumulation of Aβ due to impaired lysosomal degradation, lipid droplet formation that is exacerbated by Aβ accumulation, induction of DAM signatures, and suppression of immune response programs that render microglia less responsive to inflammatory stimuli. In parallel, SHIP1-deficient microglia exhibited NLRP3 inflammasome activation, a process we identify as being driven by lysosomal dysfunction and cathepsin B leakage. Importantly, pharmacological inhibition of cathepsin B suppressed excessive IL-1β and IL-18 release following SHIP1 inhibition or lysosomal damage, while antibody-mediated modulation of TREM2 partially rescued excessive phagocytosis and impaired lysosomal degradation in SHIP1-deficient microglia, highlighting multiple potentially targetable pathways downstream of SHIP1 deficiency. Moreover, proteomic analysis of genetically diverse human microglia demonstrated that both SHIP1 abundance and the AD-associated rs10933431 risk allele are linked to reduced abundance of SHIP1 phosphatase-domain peptides and altered endo-lysosomal programs, providing a genetic link between impaired SHIP1 function and dysfunctional microglial states in AD. Taken together, our findings point to loss of SHIP1 function as a key molecular event that drives the convergence of multiple pathogenic microglial phenotypes observed in AD.

## METHODS

### Mice

All animal procedures were conducted in accordance with the guidelines set forth by the National Institutes of Health and the University of California, Irvine Institutional Animal Care and Use Committee (IACUC; Protocol #AUP-23-074) and MGB/BWH IACUC (Protocol #2016N000464). 5x-hFIRE mice (5xFAD^+/-^, Csf1r^FIREΔ/Δ^, Rag2^-/-^, iL2rg^-/-^, hCSF1^+/+^) were previously generated and genotyped as described in Chadarevian et al^64^. Briefly, hFIRE (Csf1r^FIREΔ/Δ^, Rag2^-/-^, iL2rg^-/-^, hCSF1^+/+^)^64^ were crossed with amyloid accumulating, xenotolerant 5x-hCSF1 mice. The resulting heterozygous 5x-hFIRE (5xFAD^+/-^, Csf1r^FIREΔ/+^, Rag2^-/-^, iL2rg^-/-^, hCSF1^+/+^) females were then bred with homozygous hFIRE males to generate hFIRE and 5x-hFIRE littermates. Importantly, all experimental 5x-hFIRE mice inherited the 5xFAD transgene maternally to prevent recently reported inheritance effects on amyloid-β plaque burden in the 5xFAD mouse model^82^. All mice were sex-matched, age-matched and group-housed on a 12-h/12-h light/dark cycle with food and water ad libitum. Mice were housed with ambient temperature and humidity, and cages and bedding were changed every 1–2 w.

Synaptosome isolation was performed at BWH using 5 or 6-month-old adult wild-type C57BL/6 mice.

### Induced pluripotent stem cell (iPSC) lines

iPSC lines were utilized following IRB review and approval through MGB/BWH IRB (#2015P001676). iPSCs were generated from cryopreserved peripheral blood mononuclear cell (PBMC) samples from autopsied participants from the ROS or MAP cohort studies^16^. iPSCs were generated using Sendai reprogramming method^17^. iPSCs undergo a rigorous quality procedure that includes a sterility check, mycoplasma testing, karyotyping, and pluripotency assays performed by the New York Stem Cell Foundation (NYSCF). iPSCs were maintained using StemFlex Medium (Thermo Fisher Scientific). All cell lines were routinely tested for mycoplasma using PCR kit (MP0035-1KT) and STR profiling to prevent potential contamination or alteration to the cell lines. iPSC cell line harboring two homozygous familial Alzheimer’s disease mutations (APP^SWE^/PSEN1^M146V^;APP^SWE^/PSEN1^M146V^) and its isogenic WT control (Coriell Institute, catalog ID: AG07889) were obtained from NYSCF and have been previously described^62^.

### CRISPR targeting to generate *INPP5D* heterozygote and knockout iPSCs

We generated iPSCs heterozygous or knockout for *INPP5D* and isogenic WT lines as we previously described^11^. Briefly, guide RNAs were designed using the Broad Institute CRISPick sgRNA design tool^83,84^. The guide RNAs were ligated into plasmid backbone (pXPR_003, Addgene) and the sgRNA plasmid along with a plasmid containing Cas9 (pLX_311-Cas9, Addgene) were transfected into iPSCs using Lipofectamine 3000. Gene editing was confirmed using the GeneArt Genomic Cleavage Detection Kit and the target locus was amplified with PCR and sent for sequencing.

### sgRNA primers used for CRISPR

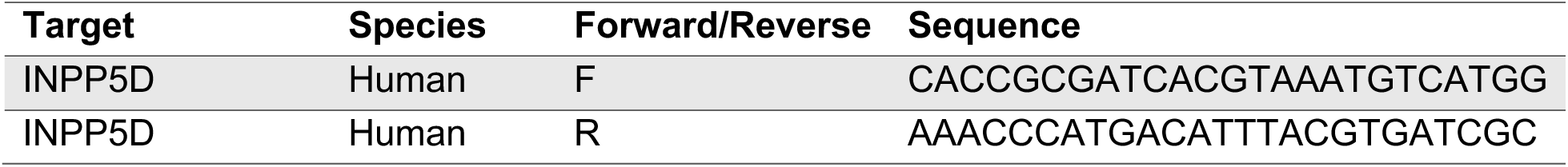

### Differentiation of iPSCs to microglia (iMGs)

iPSC-derived microglia (iMGs) were differentiated following a previously published protocol^85,86^ with minor modifications^11^. iPSCs were plated on growth factor reduced Matrigel (Corning, #354230) using StemFlex Medium (Thermo Fisher Scientific) and ROCK inhibitor (10 mM). From day 0 to day 12 of differentiation, StemDiff Hematopoietic Kit (StemCell Technologies) was used to generate hematopoietic precursor cells (HPCs). On day 12, cells were replated at 10,000 cells/cm^2^ in iMG media (DMEM/F12, 2X insulin-transferrin-selenite, 2X B27, 0.5X N2, 1X GlutaMAX, 1X non-essential amino acids, 400mM monothioglycerol, 5 mg/mL insulin, and 1% Pen-Strep) supplemented with 3 cytokines (IL-34 (100 ng/mL, PeproTech), TGF-β1 (50 ng/mL, Militenyi Biotech), and M-CSF (25ng/mL, Thermo Fisher Scientific)). From days 12–20, iMG media with freshly added cytokines was added to the culture every other day. On day 20, cells were banked in BamBanker and later thawed in fresh iMG medium at 100,000 cells/well in 24 well plates coated with growth factor reduced Matrigel. From day 20 to day 37, iMG media with freshly added 3 cytokines were added to the culture every other day. On day 37, cells were dissociated with PBS at room temperature for 5 minutes, counted, and re-plated onto final experimental plates. Starting at day 37, the iMG media was supplemented with two additional cytokines: CD200 (100 ng/mL, Novoprotein) and CX3CL1 (100 ng/mL, Peprotech) every other day until harvest at day 40.

### Differentiation of iPSCs to neurons (iNs)

iPSC-derived neurons (iNs) were differentiated following a previously published paper^87^ with minor modifications^17^ iPSCs were plated at a density of 95,000 cells/cm^2^ on plates coated with growth factor reduced Matrigel one day prior to virus transduction (Corning #354230). Then, iPSCs were transduced with three lentiviruses – pTet-O-NGN2-puro (Addgene plasmid #52047, a gift from Marius Wernig), Tet-O-FUW-EGFP (Addgene plasmid #30130, a gift from Marius Wernig), and FUdeltaGW-rtTA (Addgene plasmid #19780, a gift from Konrad Hochedlinger). The cells were then replated at 200,000 cells/cm^2^ using StemFlex Medium (Thermo Fisher Scientific) and ROCK inhibitor (10 mM) (day 0). The media was changed to KSR media (Knockout DMEM, 15% KOSR, 1x MEM-NEAA, 55 mM β-mercaptoethanol, 1x GlutaMAX (Life Technologies) on day 1, 1:1 of KSR and N2B media (DMEM/F12, 1x GlutaMAX (Life Technologies), 1x N2 supplement SB (StemCell Technologies), 0.3% dextrose (D-(+)-glucose, Sigma)) on day 2 and N2B media (Neurobasal medium, 0.5x MEM-NEAA, 1x GlutaMAX (Life Technologies), 0.3% dextrose (D-(+)-glucose, Sigma)) on day 3. On day 4, cells were dissociated using Accutase and plated at 50,000 cells/cm^2^ using iN D4 media (NBM media + B27 (1:50) + BDNF, GDNF, CNTF (10 ng/mL, PeproTech). Doxycycline (2mg/ml, Sigma) was added from day 1 to the end of the differentiation, and puromycin (5mg/ml, Gibco) was added from day 2 to the end of the differentiation. On day 3, B27 supplement (1:100) (Life Technologies) was added. From day 4 to the end of differentiation day 21, cells were cultured in iN day 4 media and fed every 2–3 days. Synaptophysin-Gamillus NGN2-iPSCs (a gift from Martin Kampmann) were differentiated into iNs without puromycin.

### iPSC-derived microglia-neuron co-culture

iNs were differentiated to day 20 and iMGs were differentiated to day 40 separately as described above. On day 40 of differentiation, iMGs were dissociated with PBS and re-plated onto iNs at 1:1 ratio of iMG:iN in BrainPhys Neuronal Medium (StemCell Technology, 5792) containing NeuroCult SM1 Neuronal Supplement (StemCell Technology, 5711), supplemented with 5 growth factors (100 ng/mL IL-34, 50 ng/mL TGF- β1, 25ng/mL M-CSF, 100 ng/mL CD200 and 100 ng/mL CX3CL1). iMGs and iNs were co-cultured for 3 days. See Lish et al. (2024) for more details^88^.

### Western blotting

Cells were lysed with RIPA lysis buffer (ThermoFisher Scientific #89900) with the protease inhibitor (Complete mini protease inhibitor, Roche) and phosphatase inhibitor (phosphoSTOP, Roche) added freshly before the lysis. Cells were lysed for 30 minutes on ice before transferring lysates to microcentrifuge tubes. Cell debris was pelleted by centrifugation (13,000 x g) for 10 minutes at 4 °C. Supernatant (cell lysate) was collected and stored at −20 °C until use. Cell lysates were prepared with 4X LI-COR loading buffer (Fisher Scientific, #NC9779096) and 2.5% β-mercaptoethanol, centrifuged, and incubated at 95 °C for 10 minutes. Samples were resolved using Novex NuPAGE 4-12% Bis-Tris gels (ThermoFisher, #NP0336BOX) and NuPAGE 1X MOPS-SDS or MES-SDS running buffer (ThermoFisher, #NP0001). Gel electrophoresis was run at 200 V for 45 minutes. SeeBlue Plus2 pre-stained protein standard (ThermoFisher, #LC5925) was used for evaluation of molecular weight. The gel was extracted and transferred to a nitrocellulose membrane by incubation with 20% methanol tris-glycine transfer buffer at 400 mA for two hours. The transferred blot was blocked with Odyssey blocking buffer (LI-COR, #927-50100) for 1 hour at room temperature with agitation and incubated with primary antibody (diluted in blocking buffer) overnight at 4 °C with agitation. Blots were incubated with LI-COR secondary antibody diluted 1:10,000 in TBST for 1 hour at room temperature with agitation. Blots were washed twice (10 minutes per wash) with TBST and stored in 1XTBS until imaging. Blots were imaged on a LI-COR Odyssey machine and quantified using ImageStudio software.

### Immunocytochemistry

Cells were washed with PBS and then fixed with 4% paraformaldehyde (PFA, Sigma) for 15 minutes at room temperature. Cells were blocked in 2% donkey serum (Jackson Immunoresearch Laboratories) and 0.3% Triton-X-100 (Sigma) in PBS for 1 hour at room temperature with agitation. Primary antibodies were diluted in a fresh donkey serum blocking buffer and cells were incubated with primary antibody solution overnight at 4 °C. Then, cells were washed with PBS three times, incubated with secondary antibodies for 1 hour at room temperature with agitation, and then washed with PBS three times. Cells were treated with DAPI stain (1:1000 dilution) for 10 minutes at room temperature with agitation, followed by a final wash to prepare for imaging. To visualize lipid droplets, BODIPY 493/503 (ThermoFisher Scientific, #D3922) or LipidSpot (Biotium, #70069) was added at 1:1000 to the staining mixture with DAPI. Images were taken on Andor Dragonfly 600 Spinning Disk Confocal at 100x or 40x magnification with oil immersion with 0.3 μm z-stacks. All images were processed in ImageJ with brightness and contrast adjustments applied consistently across all images for representative images.

### Adult intracranial transplantation of HPCs

For adult intracranial transplantation, WT or *INPP5D* HET iPSC-derived hematopoietic progenitor cells (HPCs) were bilaterally transplanted into the hippocampus and overlying cortex of 2-month-old adult hFIRE and 5x-hFIRE mice. Briefly, adult mice (2 months old) were anesthetized under continuous isoflurane and secured to a stereotaxic frame (Kopf), and local anesthetic (Lidocaine 2%) was applied to the head before exposing the skull. Using a 30-gauge needle affixed to a 10 mL syringe (Hamilton #7659-01), mice received 2 ul of HPCs in sterile 1X DPBS at 62,500 cells/uL at each injection site. Transplantation was conducted bilaterally in the cortex and hippocampus at the following coordinates relative to bregma: anteroposterior, 2.06 mm; mediolateral, ± 1.75 mm; dorsoventral, 1.75 mm (hippocampus) and 0.95 mm (cortex). Cells were injected at a rate of 62,500 cells/ 30 s with 4 min in between injections. The needle was cleaned with consecutive washes of DPBS, 70% (vol/vol) ethanol, and DPBS in between hemispheres and animals. Animals were allowed to recover on heating pads before being placed in their home cages and received 2 mg/mL Acetaminophen (Mapap) diluted in water for ten days.

### Tissue collection for IHC and proteomics

Animals were perfused at 6-months of age with 1X DPBS for 3 min at 4°C. Each mouse’s whole brain was surgically excised and cut into halves by a single mid-sagittal cut. Per mouse, one hemisphere was placed in 4% paraformaldehyde (PFA) for 24 hours at 4°C then transferred to PBS with 0.05% NaN_3_, and the other hemisphere was fresh-frozen after removal of the cerebellum and pulverized on dry ice using a mortar and pestle.

### Immunohistochemistry

Flash-frozen mouse brain hemispheres were drop-fixed in 4% PFA for 24 hours at 4 °C and subsequently transferred to PBS containing sodium azide (PBS + NaN₃) for long-term storage at 4 °C. Prior to sectioning, brains were cryoprotected in a sucrose gradient (10%, 20%, and 30% sucrose in PBS). Brains were embedded in OCT and sectioned coronally at 50 µm thickness using a cryostat. Free-floating sections were stored in cryopreservative solution (31.25% glycerol, 31.25% ethylene glycol, 37.5% PBS) at −20 °C until immunostaining.

For immunostaining, sections were washed with PBS and incubated in Liberating Antibody Binding (LAB) solution (Polysciences) for 20 minutes at room temperature with agitation. Sections were washed with PBS three times, then blocked in 2% donkey serum and 0.3% Triton-X-100 in PBS for 1 hour at room temperature with agitation. Sections were incubated with primary antibody solution for 1-2 days at 4 °C. Then, sections were washed with PBS three times, incubated with secondary antibodies for 1 hour at room temperature with agitation, and then washed with PBS three times. Sections were treated with DAPI stain (1:1000 dilution) for 10 minutes at room temperature with agitation, followed by a final wash. To visualize lipid droplets, BODIPY 493/503 (ThermoFisher Scientific, #D3922) was added at 1:1000 to the staining mixture with DAPI. Prior to mounting, sections were incubated in TrueBlack Plus (Biotium) at 1X for 10 minutes at room temperature with agitation, followed by three washes with PBS. Sections were mounted with Vectashield HardSet Antifade Mounting Medium (Vector Laboratories). High-resolution images were taken on Andor Dragonfly 600 Spinning Disk Confocal at 100x magnification with oil immersion with 1 μm z-stacks. Low-resolution images were taken on AxioScan 7 (Zeiss) at 20x magnification. All images were processed in ImageJ with brightness and contrast adjustments applied consistently across all images for representative images.

### Antibodies for western blot, immunocytochemistry and immunohistochemistry

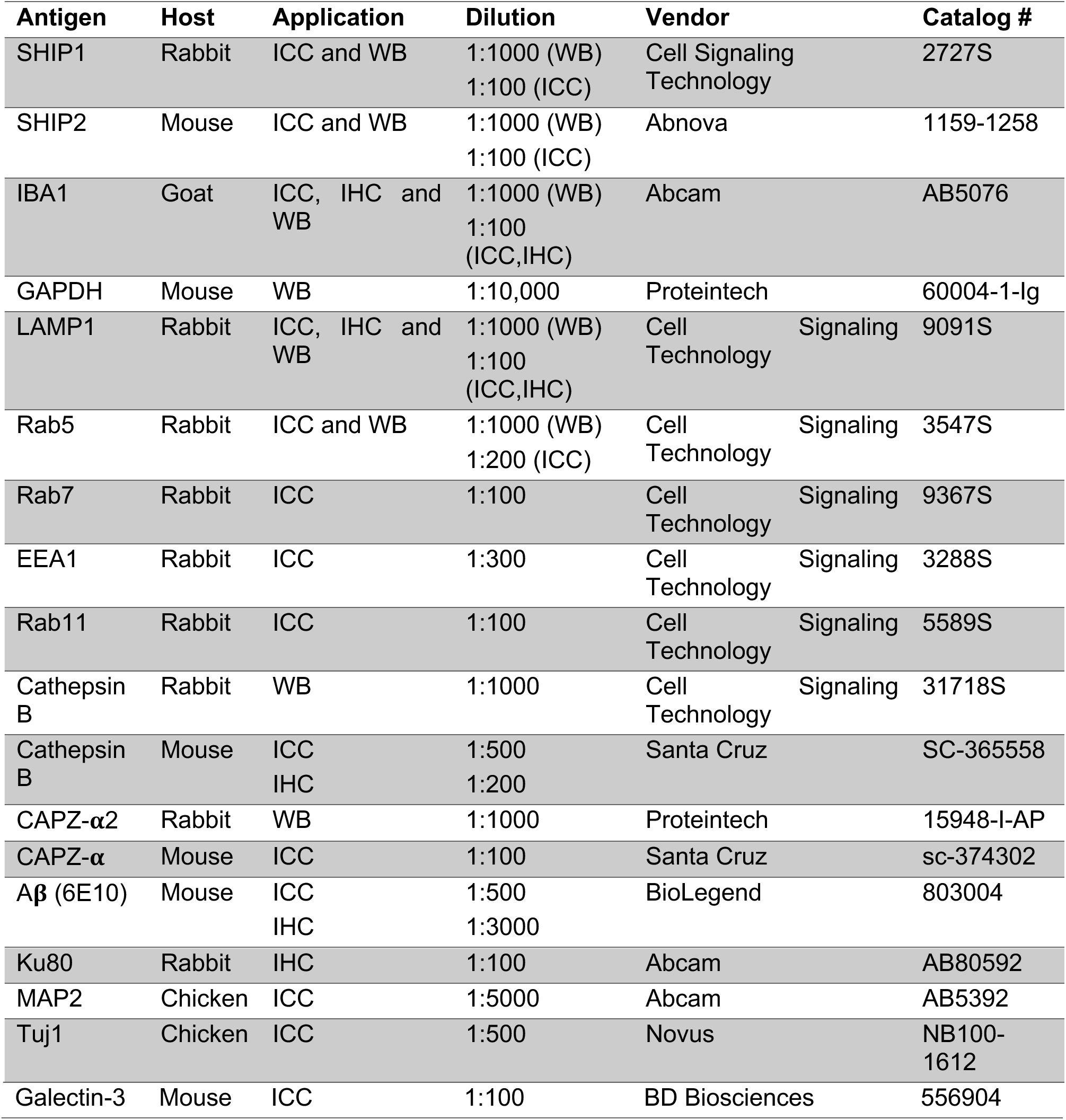

### qPCR

On D40 of differentiation, iMGs were harvested and RNA was purified using Purelink RNA Mini kit (Invitrogen). cDNA was generated using SuperScript II (Invitrogen). qPCR was performed using Power SYBR Green Master Mix and run on ViiA7 system (Applied Biosystems). Samples were assayed with 3 technical replicates and analyzed using the DDCt method and expression was normalized to GAPDH expression.

### Primers used for qPCR

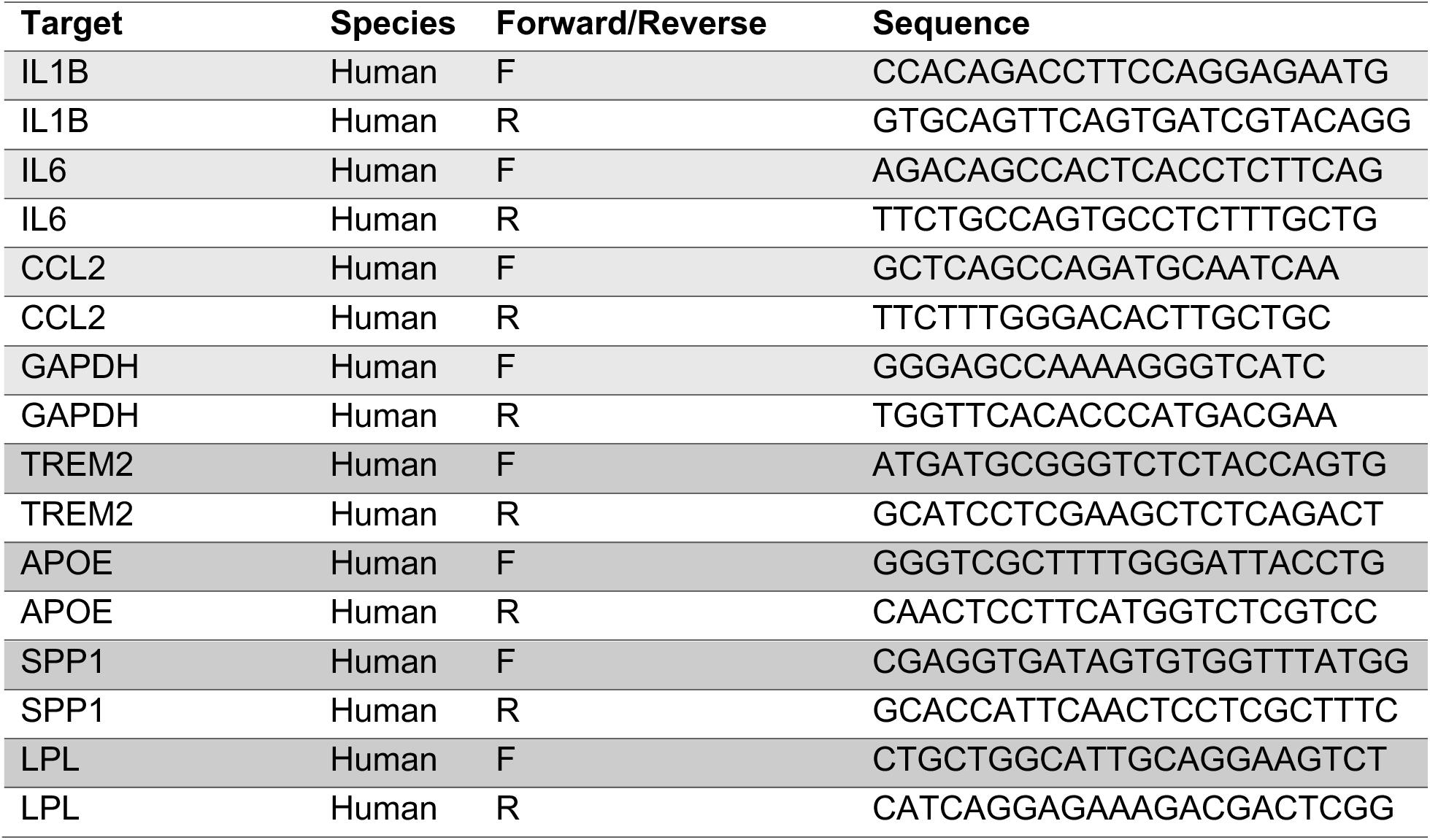

### ELISA kits used for secreted protein measurements

| Analyte(s) | Species | Vendor | MSD Assay Kit | Catalog number |
| --- | --- | --- | --- | --- |
| IL-1β, IL-18 | Human | MesoScale Discovery | U-PLEX | K15067L-2 |
| IL-6 | Human | MesoScale Discovery | V-PLEX | K151QXD-2 |
| APOE | Human | MesoScale Discovery | R-PLEX | K1512IR-2 |
| SPP1 | Human | MesoScale Discovery | R-PLEX | K151YMR-2 |
| Aβ38, Aβ40, Aβ42 | Human | MesoScale Discovery | V-PLEX | K15200E-2 |

### Immunoprecipitation followed by mass spectrometry (IP-MS)

BR33 WT and BR24 WT and *INPP5D* KO iMGs were lysed in 1% CHAPSO buffer and protein concentration was measured using Pierce BCA Protein Assay Kit (ThermoFisher, #23225). Lysates were pre-cleared with Protein G Dynabeads (Invitrogen, #10003D). SHIP1 antibody (ab45142) was cross-linked to Protein G Dynabeads with BS_3_ (ThermoFisher, #21580). 5 μg of SHIP1 antibody was used for immunoprecipitation. Immunoprecipitation was performed by incubating pre-cleared lysates with the cross-linked antibody-Dynabeads complex overnight at 4 °C. For immunoprecipitation followed by Western blotting, beads were washed 3 times with 1% CHAPSO lysis buffer before eluting protein with 5% β-mercaptoethanol and loading sample onto a Novex NuPAGE 4-12% Bis-Tris gel (ThermoFisher, #NP0336BOX). For immunoprecipitation followed by mass spectrometry, beads were washed with CHAPSO lysis buffer and frozen at −80 °C. Samples were sent to the Emory Integrated Proteomics Core (Emory University) for peptide sequencing and identification. Enrichment analysis was performed with Perseus according to guidance from the Emory Integrated Proteomics Core.

Gene ontology (GO) cellular component (CC) and Reactome pathway analyses were performed on the list of SHIP1 binding partners using DAVID (https://david.ncifcrf.gov). P-values were adjusted using Benjamini-Hochberg procedure (false discovery rate (FDR) < 0.05).

### Proximity ligation assay

Proximity ligation assay was performed according to the manufacturer’s protocol (Sigma-Aldrich, #DUO82049). Briefly, D40 iMGs were fixed with 4% paraformaldehyde at room temperature for 15 minutes and permeabilized with 0.2% Triton-X 100 in PBS for 1 hour. After permeabilization, cells were blocked with Duolink blocking solution for 1 hour 37 °C. Primary antibodies were diluted to 1:500 in Duolink Antibody Diluent and incubated for 1 hour at room temperature. The samples were washed twice with Wash buffer A for 5 minutes. Secondary probes were diluted 1:7 in Duolink Antibody Diluent and incubated for 1 hour at room temperature. The samples were washed twice with Wash buffer A for 5 minutes. Immediately before use, the ligase was diluted 1:80 in 1X ligation buffer and incubated for 30 minutes at 37 °C. The samples were washed twice with Wash buffer A for 5 minutes. DNA polymerase was diluted 1:80 in 1X amplification buffer immediately before use and incubated for 30 minutes at 37 °C. The samples were washed twice with Wash Buffer B for 10 minutes and an additional minute with 0.1X Wash Buffer B.

For additional immunocytochemistry, secondary antibodies were diluted in Duolink Antibody Diluent at 1:2000 and incubated for 1 hour at room temperature. The samples were washed twice with Wash Buffer B for 10 minutes and an additional minute with 0.1X Wash Buffer B. Samples were mounted with DAPI mounting solution overnight at room temperature.

Four types of controls were included in each experiment to validate specificity. For the single target primary negative control, only one primary antibody was used, along with both PLA secondary antibodies (from different species) bearing complementary oligonucleotide strands. For the single target secondary control, both primary antibodies were added, but only one PLA secondary antibody was included. For the genetic knockout, primary and secondary antibodies were included, but the experiment was performed in *INPP5D* KO iMGs. For the positive control, a single primary antibody was paired with two PLA secondary antibodies targeting the same species, each carrying complementary oligonucleotide strands. These controls were used to ensure the observed PLA signal reflected specific protein-protein proximity rather than nonspecific interactions or background.

### Endo-lysosomal measurements

#### DQ-Red BSA Assay

To measure lysosomal degradation activity, iMGs were incubated in 1 μg/mL DQ-Red BSA (Invitrogen, #D12051) for 1 hour at 37 °C, after which the wells were washed once with PBS and fresh iMG media was added for the 2-hour chase period. 9 images per well for each well were acquired at 20x at the end of the chase period using IncuCyte S3 Live-Cell Imaging System (Sartorius). Average DQ-BSA intensity was calculated by dividing the integrated DQ-Red BSA intensity by microglia area, measured by the IncuCyte S3 software (Sartorius). For negative control wells, iMGs were pre-treated with 100 nM bafilomycin A1 (Tocris Biosciences, #1334) for 6 hours prior to addition of DQ-Red BSA. The cells were fixed with 4% PFA for 15 minutes at room temperature at the end of imaging for immunostaining.

#### BSA endocytosis assay

To measure endocytosis of BSA, iMGs were incubated with 1 μg/mL BSA-Alexa 488 (Invitrogen, Cat. No. A13100) for 1 hour at 37 °C. Cells were dissociated with PBS at room temperature for 5 minutes, after which they were incubated in Zombie Violet (BioLegend, #423113) at 1:1000 dilution for 15 minutes on ice. The cells were washed once with PBS before fixation 4% PFA for 15 minutes on ice. The cells were washed again with PBS and suspended in FACS buffer for flow cytometry measurement of BSA-Alexa 488 fluorescence. Negative control wells were pre-treated with 10 μM cytochalasin D (Sigma-Aldrich, #C8273) for 30 minutes prior to the addition of BSA-Alexa 488.

#### LysoTracker

For LysoTracker measurements, cells were incubated with 50 nM LysoTracker Green (Invitrogen, #L7526) for 1 hour at 37 °C. Cells were dissociated with room temperature PBS for 5 minutes, spun down at 500 g for 5 minutes, then incubated in Zombie Violet (BioLegend, 1:1000) for 15 minutes on ice. The cells were spun down again and resuspended in FACS buffer for flow cytometry measurement of LysoTracker fluorescence.

#### pHrodo-dextran assay

To measure endosome acidification, Zombie Violet was diluted in the media in iMG wells at 1:1000 and cells were incubated for 5 minutes at 37 °C. After 5 minutes, pHrodo-dextran (Invitrogen, #P35368) was added to the wells at 50 μg/mL and cells were incubated for another 5 minutes at 37 °C. The media with Zombie Violet and pHrodo-dextran was then removed and replaced with fresh iMG media. The cells were either collected after 15 minutes in fresh iMG media or at the end of the pHrodo-dextran incubation for baseline measurement. Cells were dissociated with PBS and immediately placed on ice for flow cytometry measurement of pHrodo-dextran intensity.

#### FIRE-pHLy

To measure lysosomal pH, iMGs were transduced with lentivirus encoding FIRE-pHLy (Addgene, #170775, MOI=10) co-delivered with Vpx VLPs as previously described^89^. The cells were transduced on D41 or D42 of differentiation and live-imaged 5 days after transduction. Cells were then fixed with 2% PFA for 15 minutes at room temperature covered from light. Cells were placed in PBS for imaging after fixation.

### Colocalization analysis for SHIP1 and DQ-BSA

Surface renderings of IBA1, DQ-BSA, and SHIP1 were generated in Imaris (Oxford Instruments) from images acquired at 100x magnification with z-stacks collected at 0.3 µm intervals. Image processing settings were standardized across genotypes and differentiations, while surface parameters were optimized for each differentiation. Colocalization analyses were then performed by first filtering puncta for complete (100%) localization within IBA1 surfaces, followed by a filter requiring ≥10% overlap between SHIP1 and DQ-BSA.

### Measurement of cathepsin B activity

Cathepsin B activity was measured using the Magic Red Cathepsin B Assay Kit (Abcam, #ab270772) following manufacturer’s instructions. Briefly, 150X staining solution was diluted 1:10 in water, and the diluted solution was added at 1:15 to the cells. iMGs were incubated for 30 minutes at 37 °C protected from light. Cells were washed twice with PBS and placed in fresh iMG media. Cells were imaged with the IncuCyte S3 Live-Cell Imaging System (Sartorius) for 18 hours and average Magic Red intensity quantified using IncuCyte S3 software.

### Inhibition of SHIP1 and cathepsin B activity

SHIP1 activity in iMGs was inhibited using different concentrations of 3AC (Echelon Biosciences, #NC1475141) for 6 hours on D40 or D41 of iMG differentiation. To inhibit the activity of cathepsin B in addition to SHIP1, iMGs were pre-treated with different concentrations of CA-074-Me (MilliporeSigma, #20-553-11MG) or equal volume of DMSO prior to the addition of 3AC or equal volume of 100% EtOH for 6 hours.

### Live imaging of LysoTracker and Magic Red

iMGs were stained with 1X Magic Red staining solution, 1X CellTracker Blue CMAC (Invitrogen, #C2110) and 50 nM LysoTracker Green for 30 minutes at 37 °C protected from light. Cells were washed twice with PBS, placed in fresh iMG media and incubated at 37 °C for 1 hour for the Magic Red signal to develop prior to imaging.

### CITE-Seq

#### Reconstitution of customized microglia-specific CITE-Seq panel

Reconstitution of the CITE-Seq/TotalSeq-A panel was performed according to the manufacturer’s instructions and as previously described^48^. Briefly, both lyophilized panels, the custom TotalSeq A (BioLegend, microglia-specific, described in detail in ^48^) and the TotalSeq-A Universal Cocktail (BioLegend, #399907) were equilibrated to room temperature for 5 minutes. Then, both panels were centrifuged at 10,000 x g for 30 seconds at room temperature and only the lyophilized custom panel was reconstituted in 27.5 μL of Cell Staining Buffer (BioLegend, #420201), vortexed for 10 seconds and incubated at room temperature for 5 minutes. After vortexing, the resuspended cocktail was centrifuged for at 10,000 x g for 30 seconds at room temperature, the entire volume (27.5 μL) of reconstituted custom cocktail was transferred to the TotalSeq-A Universal Cocktail vial, vortexed and incubated at room temperature for 5 minutes. Following vortexing and centrifugation at 10,000 x g for 30 seconds at room temperature, the entire volume (27.5μL) of reconstituted combined cocktail was transferred to a low protein binding Eppendorf tube (Thermo Fisher Scientific, #022431081) and centrifuged at 14,000 x g for 10 min at 4 °C. After this step, the cocktail was immediately added to the cell preparation following 5 minutes of incubation in Human TruStain FcXTM Fc blocking reagent (BioLegend, #422301).

#### CITE-Seq staining of iMGs

CITE-Seq staining protocol was performed as previously described^48^. Briefly, D40 BR33 WT and HET iMGs with technical duplicates were dissociated with PBS for 5 minutes at room temperature, collected, counted and cell viability was assessed. 400,000 cells per condition were transferred to 5 mL low-protein binding tubes (Thermo Fisher Scientific, #13-864-407). 1 mL cold DPBS was added to the tubes, after which the cell-DPBS mixture was taken up and filtered through a blue lid filter cap back into the tubes. The tubes were spun down at 300 x g for 10 minutes at 4 °C, followed by repeating of the washing step. The supernatant was carefully removed, and the cell pellet was resuspended in 22.5 μL of Cell Staining Buffer (BioLegend, #420201). 2.5 μL of Human FcBlock (BioLegend, #422301) was added and cells were incubated for 10 minutes on ice. Then, 6.25 μL of reconstituted combined CITE-Seq cocktail was mixed with 18.75 μL Cell Staining Buffer, added to the cells, and incubated for 30 minutes on ice. Then, cells were washed with 1 mL Cell Staining Buffer, spun down at 300 x g for 10 minutes at 4 °C, followed by repeating of the washing step 3 more times. Before the last wash, cells were filtered through a blue lid filter cap back into the tubes. Cells were counted and processed for cDNA library preparation.

#### Library preparation

After washing, the blocked and surface-stained cells were filtered through 30µm Celltrics filters, counted and volume adjusted to bring cell concentrations to 4,000 cell/µl. Gel emulsions were generated using the 10xGenomics Chromium controller and NextGEM 3’ reagents v3.1 (10x Genomics), using 16µls of prepared sample to target 64,000 loaded cells into 4 wells of GEM G chip. Emulsions were uniform and fully formed for library preparation. Libraries were generated according to the Chromium Next GEM Single Cell 3’ Reagent Kits v3.1 (Dual Index) user guide CG000317 Rev D. After generation of bar-coded single-cell library pools, cDNAs were cleaned and amplified. The amplified pools were size selected and split into pools for generating either gene expression (GEX) or surface protein (PEX) libraries according to user guide CG000317. The final amplified and indexed libraries demonstrated expected size distributions using a TapeStation 4200 to assay quality. All gene expression libraries ranged from 350-650bp and surface protein libraries all contained a single band of ∼225bp. GEX and PEX libraries were independently normalized and pooled and sequenced on a NovaSeq 6000 using an S4 flow cell to a targeted depth of 20,000 PE reads for GEX and 10,000 PE reads for PEX libraries per cell.

#### Data preprocessing

Sequencing output was processed using the Cell Ranger pipeline from 10x Genomics v7.2.0 in a computing cluster environment using the GRCh38-2020-A human reference genome. A custom feature mapping file was used to assign identified antibody tags to a cell surface protein count matrix. CellRanger output was then imported to an R environment on a stand-alone machine to run QC and test differential expression. We used the Seurat package (v5.3.0) for QC, normalization, clustering, and differential expression analyses. Cells with greater than 10,000 UMIs detected, and between 500 and 4,000 detected RNA features, and fewer than 15% mitochondrial genes were retained for downstream analyses with all other cells being discarded. For the ADT assay, cells with 50–105 detected features and total ADT counts between 200 and 3,000 were included, with all other cells excluded from further analysis. Merged and filtered gene expression data set was normalized using the SCTransform function in Seurat while surface protein data were normalized using the ‘CLR’ method to center-scale values. The remaining dataset had 107,256 cells (30,933 WT iMGs, 76,323 cells HET iMGs) with an average of 2284 mapped genes detected per cell and 4961 average total unique molecular identifiers (UMIs) per cell. RNA and ADT modalities were then integrated using weighted nearest neighbor (WNN) analysis and a multimodal UMAP embedding was generated. Final clustering was performed on the WNN graph using the Leiden algorithm (resolution = 0.5) to define multimodal cell populations. Cluster-specific marker genes and proteins were identified with FindAllMarkers in Seurat (v5.3.0), using the Wilcoxon rank-sum test. The results of these analyses can be found in Supplementary Tables 4 and 5.

Differential gene expression across genotypes within individual clusters was calculated by subsetting the Seurat object into individual clusters, followed by running FindMarkers to identify positively and negatively enriched markers of HET iMGs.

#### Gene ontology (GO) enrichment analysis for RNA markers

Gene ontology (GO) enrichment analysis was performed using the enrichGO function under the clusterProfiler package in R with no nominal p-value cut-off. Only biological process (BP) and molecular function (MF) terms were included in the analysis. The full results of GO enrichment analysis can be found in Supplementary Table 6.

### Synaptosome isolation and pHrodo conjugation

Synaptosomes were isolated from mouse brains using Syn-PER (Thermo Fisher Scientific, # 87793), following the manufacturer’s protocol and as previously described^89,90^ with minor modifications. Briefly, phosphatase and protease inhibitors were added to Syn-PER at a concentration of 1 tablet per 10 mL. Mouse brain tissue was weighed, and 10 mL of Syn-PER was added per gram of tissue. The tissue was homogenized on ice using a Dounce homogenizer with ∼12 slow strokes. The homogenate was centrifuged at 1200 x g for 10 minutes at 4 °C to remove debris, and the supernatant was collected. This was further centrifuged. at 15,000 x g for 20 minutes at 4 °C to pellet the synaptosomes. The final pellet was resuspended in 0.1 M sodium bicarbonate buffer (pH 8.3–8.4) at a concentration of 50 mg/mL. For pHrodo conjugation, the synaptosome suspension was incubated with 3 mM of amine-reactive pHrodo Green STP dye (Thermo Fisher Scientific) for 45 minutes at room temperature in the dark. The mixture was diluted 1:10 in DPBS and centrifuged at 2500 x g for 5 minutes at 4 °C. The pellet was washed twice with DPBS and resuspended in phenol-red free DMEM/F12 to a final concentration of 50 mg/mL. Synaptosomes were used for treatments at a final concentration of 1 mg/mL.

### Apoptotic neuron preparation

Apoptotic neurons (ANs) were generated from D14 iNs as previously described^89^. Briefly, iNs were subjected to ultraviolet radiation using an ultraviolet cross-linker at 500 Jm^−2^ and collected in PBS 24 hours later. ANs were pelleted by centrifugation at 300 x g for 5 minutes. The pellet was weighed and resuspended in 0.1 M sodium bicarbonate at 50 mg/mL and conjugated with pHrodo as described in “*Synaptosome isolation and pHrodo conjugation*”. Following PBS washes, pHrodo-conjugated Ans were resuspended in 1 mL phenol-red free DMEM/F12, counted and added onto D40 iMGs at 35,000 ANs/cm^2^.

### Fibrillar Aβ generation

FITC-labeled Aβ1-42 monomers (Anaspec) were dissolved in 7 M guanidine hydrochloride overnight at room temperature. The solution was chromatographed on a Superdex 75 Increase column in running buffer 50 mM ammonium bicarbonate pH 8.5, and a single monomer peak was collected, pooled, diluted to 10 μM in TBS (20 M Tris, 500 mM NaCl, pH 7.4) using UV absorbance with extinction coefficient 1,490 M^-1^cm^-1^. To generate fibrils, 0.5 ml SEC-purified monomer was shaken at 800 rpm in a microcentrifuge tube in an Eppendorf tube-shaker at 37 ^◦^C overnight. The suspension was then centrifuged for 1 hour at 100,000 xg in a TLA-55 rotor, followed by two washes in cold TBS with the same centrifugation. The final pellet was resuspended to a concentration of 100 μM followed by sonication five times for 5 s at 35% power to disperse. The fibrils were aliquoted and flash-frozen at −80^◦^C.

### Aβ degradation assay

iMGs were treated with 0.1 μM FITC-labeled fibrillar Aβ for 30 minutes. To quantify baseline Aβ uptake, Zombie Violet was spiked into the wells at 1:1000 for the last 10 minutes of Aβ, after which the cells were dissociated with PBS and fixed with 4% PFA for 10 minutes. FITC (Aβ) fluorescence within iMGs was measured via flow cytometry.

To investigate the degradation of iMGs over 24 hours, iMGs were washed with PBS and placed in fresh iMG media after 30 minutes of Aβ. Cells were imaged every two hours using the IncuCyte S3 live-imaging system for 24 hours. After 24 hours, iMGs were dissociated with PBS, incubated in 1:1000 Zombie Violet for 10 minutes and fixed with 4% PFA for 10 minutes. FITC (Aβ) fluorescence remaining within iMGs was measured via flow cytometry.

### Phagocytosis assay

D37 iMGs were replated at 30,000 cells/well onto 48 well plates. Prior to the addition of synaptosomes, ANs or Aβ, iMGs were treated separately with the following compounds: 10 μM cytochalasin D (Sigma-Aldrich, #C8273) for 30 minutes, 10 µg/mL TREM2 IgG (R&D Systems, #AF18281) for 5 minutes, 10 µg/mL Goat IgG (R&D Systems, #AB108C) for 5 minutes and 100 nM bafilomycin A1 (Tocris Biosciences, #1334) or equal volume DMSO for 1 hour. Cells were then exposed to pHrodo-conjugated synaptosomes, ANs or FITC-labeled Aβ overnight. iMGs were gently dissociated with PBS for 5 minutes and centrifuged at 500 x g for 5 minutes at 4°C. Cells were stained with Zombie Violet (BioLegend) for live/dead discrimination and incubated on ice for 10 minutes, protected from light. After washing and centrifugation, cells were stained with PE-conjugated anti-P2RY12 and PE-Cy7-conjugated anti-CD11b antibodies in FACS buffer (HBSS supplemented with 10% FBS, 25 mM HEPES, and 2 mM EDTA). Staining was performed for 30 minutes on ice, followed by fixation with 4% paraformaldehyde for 10 minutes. Samples were analyzed on a Fortessa flow cytometer equipped with 355 nm, 488 nm, 561 nm, and 640 nm lasers. Phagocytosis was quantified as the percentage of pHrodo-positive cells within live, P2RY12+ and CD11b+ gated populations. Data acquisition and analysis were performed using FlowJo software, with compensation applied for multicolor fluorescence detection.

### Flow cytometry measurement of surface protein expression

D40 iMGs were gently dissociated with PBS for 5 minutes and centrifuged at 500 x g for 5 minutes at 4°C. Cells were stained with Zombie Violet (BioLegend) for live/dead discrimination and incubated on ice for 10 minutes, protected from light. After washing and centrifugation, cells were stained with PE-conjugated anti-TREM2 or PE-conjugated anti-CD172a/b (SIRP-α) and PE-Cy7-conjugated anti-CD11b antibodies in FACS buffer (HBSS supplemented with 10% FBS, 25 mM HEPES, and 2 mM EDTA). Staining was performed for 30 minutes on ice, followed by fixation with 4% paraformaldehyde for 10 minutes. Samples were analyzed on a Fortessa flow cytometer equipped with 355 nm, 488 nm, 561 nm, and 640 nm lasers.

### Antibodies for flow cytometry

| Antigen | Host | Dilution | Vendor | Catalog # |
| --- | --- | --- | --- | --- |
| CD11b | Rat | 1:200 | BioLegend | 101215 |
| P2RY12 | Mouse | 1:20 | BioLegend | 392104 |
| TREM2 | Rat | 1:50 | R&D Systems | FAB17291P |

### Analysis of dendritic spines

Dendritic spines in iN-iMG co-cultures were labeled using the fluorescent dye 1,1′-dioctadecyl-3,3,3′,3′-tetramethylindocarbocyanine perchlorate (DiI) by adapting a previous protocol^91^. Briefly, PBS was completely removed from each well, and forceps were used to obtain and sprinkle 3-5 DiI crystals across each well. A small amount (∼10uL) of PBS was added to the edge of each well to prevent iNs from drying and peeling away. Plates with DiI crystals were incubated on an orbital shaker for 10 minutes at room temperature. Following incubation, wells were washed 2-3 times with PBS to wash away DiI crystals. PBS was then added to each well and plates were covered to protect from light and placed at 4°C overnight, after which the wells were washed 3 times with water at room temperature for 5-10 minutes. PBS was added back into each well, plates were covered to protect from light and allowed to incubate at 4°C for at least 72 hours prior to imaging to allow for the dye to completely incorporate into the cells. Five dendrite images per well were collected using a 100X (oil immersion) objective of Dragonfly 600 spinning disk confocal microscope with 0.2 µm z-steps. Images were imported into NeuronStudio^92^ for analysis of dendritic spine density. Spine density measurements were acquired by reconstructing the dendritic cable to acquire the length of each imaged dendrite, followed by manual identification of spines for each dendrite.

### Collection and analysis of iMG images

iMG images were taken using the 100X or 40X (both oil immersion) objectives of Andor Dragonfly 600 spinning disk confocal microscope with 0.3 µm z-steps. ImageJ was used to generate maximum-intensity projections of individual channels. CellProfiler was used to establish automated pipelines to measure signal area and intensity in confocal images. IdentifyPrimaryObjects was used to segment proteins of interest, IBA1^+^ microglia and DAPI^+^ nuclei. RelateObjects was used to segment microglia containing nuclei and exclude debris. MaskObjects was used to remove any signal outside of the microglia bounds. MeasureImageAreaOccupied was used to measure area occupied by the proteins of interest and the microglia, and MeasureObjectIntensity was used to quantify signal intensity.

### Data-Independent Acquisition Mass Spectrometry (DIA-MS) proteomics of iMGs from ROSMAP donors

#### Proteomics sample pre-processing

75 independent iPSC-derived microglial lines (iMGs) generated from PBMCs collected from participants in the ROS, MAP, MARS, and AA CORE study cohorts were differentiated as described above, with cell pellets harvested at day 40 of differentiation and subsequently flash frozen at −80°C. Samples were sent to the Emory Integrated Proteomics Core (Emory University) for peptide sequencing and identification. Cell pellets were lysed with 100µL EasyPep lysis buffer (ThermoFisher, #A45735), and a Bicinchoninic Acid (BCA) assay was performed to determine protein concentration. Sample volumes equivalent to 15 µg were obtained and reduced with 10 mM Tris(2-carboxyethyl)phosphine (TCEP) and 40 mM 2-chloroacetamide (CAA) at 90°C for 10 min in the dark. Samples were digested with 1µg Lysl endopeptidase (Wako, #125-05061) and 1 µg trypsin (ThermoFisher, #90057) overnight at room temperature. Digestion mixtures were cleaned with Water’s 10mg MCX plates (Fisher, #50-780-153) according to manufacturer’s protocol, and eluents dried under speed vacuum. Each sample was resuspended in 20 µL of loading buffer (0.1% formic acid) and 1 µL was analyzed by liquid chromatography coupled to tandem mass spectrometry using a Data-Independent Acquisition Mass Spectrometry (DIA-MS) proteomics approach. Peptide eluents were separated by a Vanquish Neo (ThermoFisher) on a 15 cm x 150 µM internal diameter (ID) silica column packed with 1.5 µm resin (Dr. Maisch). Buffer A was water with 0.1% (vol/vol) formic acid, and buffer B was 100% (vol/vol) acetonitrile in water with 0.1% (vol/vol) formic acid. Elution was performed over a 14-minute gradient ranging from 3% to 50% solvent B. Peptides were monitored on an Orbitrap Astral mass spectrometer (ThermoFisher) fitted with a high-field asymmetric waveform ion mobility spectrometry (FAIMS Pro) ion mobility source (ThermoFisher). One compensation voltage (CV) of −35 was chosen for the FAIMS. Each cycle consisted of one full scan (MS1) with an m/z range of 380-980 at 240,000 resolution, 500% Automatic Gain Control (AGC) and 3 millisecond injection time. The higher energy collision-induced dissociation (HCD) data-independent acquisition (DIA) scans (MS2) were collected with a 2 m/z isolation window over the entire precursor range (380-980 m/z) with a time of 0.6 seconds and 3 millisecond injection time. Collision energy was set to 25% and scan range set to 150-2000 m/z.

#### Proteomics data analysis

Quantitative protein-level and peptide-level report measurements were acquired from the fragment ion MS2 spectra and aligned to known proteomic sequences using Spectronaut DIA software (Biognosys), yielding 132,067 peptide entries across 10,295 protein groups quantified. Both proteome-level and peptide-level data were first filtered for low-abundance proteins that were not present in more than 25% of the samples. The remaining missing values in the retained matrices were imputed using *K*-nearest neighbors (*K*NN; *k* = 10, rowmax = 0.25, colmax = 0.20, seed = 123) using the impute::impute.knn() function in R. Both protein-level and peptide-level data were mean-centered by dividing each value by the protein/peptide’s cross-sample mean. Finally, data were plating batch-corrected using the sva::ComBat() ComBat algorithm^93^, using an intercept-only (blind) model. All proteome-level abundance visualizations, correlations, and GSEA/GSVA strategies were performed in an R Bioconductor environment. Peptide-level abundance visualizations were processed using R or GraphPad Prism 10. Proteomics raw data and Spectronaut processed protein-level and peptide-level reports are available at Synapse (www.synapse.org) under the accession ID syn71985384.

#### INPP5D correlation analysis

To identify proteomic programs associated with SHIP1 abundance, pairwise correlations were calculated between SHIP1 protein abundance and the abundance of all other detected proteins across iMG samples. Protein abundances were log2-transformed prior to analysis. Pearson correlation coefficients were calculated using the cor.test function in R. Correlation results were ranked according to Pearson correlation coefficient, and positively and negatively correlated proteins were used for downstream gene set enrichment analyses.

#### GSVA analysis of lysosomal proteins

To quantify lysosomal protein abundance across iMGs, we generated a lysosome protein signature from the leading-edge genes contributing to enrichment of the KEGG Lysosome pathway identified in GSEA. Protein abundance values were averaged across technical replicates for each donor and gene symbol. A protein-by-sample abundance matrix was generated with gene symbols as rows and donor identifiers as columns. Gene Set Variation Analysis (GSVA) was performed using the GSVA R package to calculate a sample-level enrichment score for the lysosome signature. GSVA was run using a Gaussian kernel on log-transformed protein abundance values, generating a relative lysosome signature score for each donor-derived iMG sample.

To assess the relationship between SHIP1 abundance and lysosomal signature abundance, SHIP1 protein levels were extracted for each donor and averaged across technical replicates. Pearson correlation analysis was then performed between SHIP1 abundance and lysosome GSVA scores across donors. Correlation coefficients (r) and nominal P-values were calculated using the cor.test function in R.

#### rs10933431 allele-dosage analysis

To identify proteomic changes associated with the AD-associated *INPP5D* SNP rs10933431, donor-derived induced microglia (iMGs) were stratified according to rs10933431 genotype and encoded as an additive dosage variable corresponding to the number of protective G alleles (C/C = 0, G/C = 1, G/G = 2). Protein abundances were averaged across technical replicates for each donor and protein. Proteins detected in at least 50% of samples were retained for downstream analysis. Dose-dependent protein abundance changes were assessed using linear modeling implemented in the limma package (v3.64.3). For each protein, abundance was modeled as a function of rs10933431 dosage while adjusting for sex and genetic ancestry.

To identify biological pathways associated with rs10933431 dosage, proteins were ranked according to both the direction and significance of the dosage effect. Ranked gene lists were analyzed using Gene Set Enrichment Analysis (GSEA) implemented in clusterProfiler.

#### SHIP1 peptide-level analysis

SHIP1 peptide abundances were averaged across technical replicates for each donor. For visualization of genotype-dependent peptide abundance patterns, peptide abundances were aggregated by donor and z-score normalized across samples. Two-sided Wilcoxon rank-sum tests were used to compare peptide abundances across G/G and C/C carriers.

To assess the distribution of detected peptides across the SHIP1 protein, peptide sequences were mapped to amino acid positions within the canonical SHIP1 protein sequence. Protein domain annotations were obtained from InterPro and Pfam databases using the InterPro REST API. Peptides were annotated according to overlap with known protein domains.

### DIA-MS proteomics of xenografted mouse brains

#### Protein Digestion

Cryopulverized mouse brains were lysed with 50 uL of 8 M urea with protease and phosphatase inhibitors (ThermoFisher) and ice batch sonicated for 10 mins. EasyPep lysis buffer (50uL) was added, and cup horn sonicated in chilled water. BCA was performed and 5 ug was used for digestion. Samples were reduced with DTT (5 mM) and alkylated with IAA (10mM) for 10 mins at 80°C. The samples were then digested with 1 ug of Lysyl endopeptidase (Wako) and 2 ug trypsin (ThermoFisher) overnight at room temperature. The peptide solutions were then acidified to 1:1 with 4% H3PO4 and cleaned with microelute MCX columns according to manufacturer’s protocol. The eluates were then dried to completeness using a SpeedVac (LabConco).

#### Liquid Chromatography and Mass spectrometry

Each sample was resuspended in 100 uL of loading buffer (0.1% FA) and 20 uL was loaded onto Evotips and analyzed by liquid chromatography coupled to tandem mass spectrometry. Peptide eluents were separated on IonOptick’s column (15 cm × 75 μM internal diameter (ID) packed with 1.7um resin) by a Evosep One (Evosep). Buffer A was water with 0.1% (vol/vol) formic acid, and buffer B was 100% (vol/vol) acetonitrile in water with 0.1% (vol/vol) formic acid. Elution was performed using the preset 40SPD Whisper Zoom method. Peptides were monitored on a Orbitrap Astral mass spectrometer (ThermoFisher) fitted with a high-field asymmetric waveform ion mobility spectrometry (FAIMS Pro) ion mobility source (ThermoFisher). One compensation voltages (CV) of −40 was chosen for the FAIMS. Each cycle consisted of one full scan (MS1) was performed with an m/z range of 380-980 at 240,000 resolution, 500% AGC and 3 ms injection time. The higher energy collision-induced dissociation (HCD) DIA scans were collected with a 3 m/z isolation windows over the entire precursor range (380-980 m/z) with a time of 0.6 seconds and 3 ms injection time. Collision energy was set to 27% and scan range set to 150 - 2000 m/z. Database Search Spectronaut (version 21.0.260602.94842) was used to search all raw files in default library free mode using a human UniProt database or a combined mouse and human UniProt database. All parameters were kept at default. Spectronaut processed protein-level reports are available at Synapse (www.synapse.org) under the accession ID syn75612435.

#### Proteomics data analysis

Proteins were classified as human-specific, mouse-specific, shared human/mouse, or other based on FASTA header annotations. Downstream analyses focused on human-specific proteins to assess proteomic changes in xenografted human microglia. Log2 transformed protein abundance values acquired from the fragment ion MS2 spectra of DIA-MS were used for analysis. Samples were assigned to four experimental groups based on mouse genotype and transplanted iMG genotype: WT-hFIRE engrafted with WT iMGs (WT-WT), WT-hFIRE engrafted with HET iMGs (WT-HET), 5x-hFIRE engrafted with WT iMGs (5xFAD-WT), and 5x-hFIRE engrafted with HET iMGs (5xFAD-HET). Proteins were retained for analysis if they were detected in at least five samples per experimental group.

Differential abundance analysis was performed using linear modeling implemented in the limma package (v3.64.3). A model matrix without intercept was constructed using the four experimental groups, and empirical Bayes moderated statistics were calculated for the following contrasts: HET versus WT iMGs within WT-hFIRE brains, HET versus WT iMGs within 5x-hFIRE brains, 5x-hFIRE versus WT-hFIRE environment within WT iMG-engrafted brains, 5x-hFIRE versus WT-hFIRE environment within HET iMG-engrafted brains, and the interaction between iMG genotype and mouse brain environment.

For pathway analysis, proteins were ranked by moderated t-statistic from limma differential abundance results.

#### INPP5D correlation analysis

To identify molecular programs associated with SHIP1 abundance in xenografted human microglia, Pearson correlation analyses were performed between human SHIP1 protein abundance and all quantified human proteins within WT-hFIRE and 5x-hFIRE cohorts separately. To identify biological pathways associated with SHIP1 abundance, proteins were ranked according to a correlation-based score defined as sign(correlation coefficient) x −log10(p-value).

### Statistical Analysis

Information regarding statistical analyses can be found in the figure legends. All statistical tests were performed using GraphPad Prism 10. All data is shown as mean ± SEM and normalized to WT and/or vehicle-treated conditions, unless otherwise stated. Statistical analysis for cell culture experiments comparing two groups across multiple differentiations were performed using a two-way mixed-effects analysis in GraphPad Prism with multiple comparisons. Data were arranged in a grouped format, with culture type (e.g., WT vs HET) as the between-subject factor (columns) and differentiation as the within-subject factor (rows). A mixed-effects model was selected under “Repeated Measures,” and a full model was fit (column/culture type effect, row/differentiation effect, and column/culture x row/differentiation interaction). The asterisks presented in bar graphs represent the statistical significance of the column/culture type effect. One-way ANOVA was used for experiments involving more than two groups, and statistical differences between the groups were calculated using Tukey’s post-hoc comparisons test. Two-way ANOVA was used for experiments involving two independent categorical variables (e.g.: WT vs HET and vehicle vs treatment), using averages from each differentiation for each condition.

### Data visualization

Schematics were generated using BioRender.com. Graphs and heatmaps were generated using R Studio or GraphPad Prism 10. The smaller data points on the graphs represent technical replicates, the larger data points represent experiment averages. The differentiations are distinguished by different shapes.

## Supporting information

Description of supplementary files

## Acknowledgments

We thank the NeuroTechnology Studio and the ARCND Flow Cytometry Core Facility at Brigham and Women’s Hospital for access to equipment and consultation, Duc Duong and the Proteomics Core at Emory University for IP-MS analysis and consultation, Andrew Stern for fibrillar Aβ generation, Martin Kampmann for providing the synaptophysin-Gamillus line, Aimee Kao for guidance on the FIRE-pHly construct and Isaac Chiu, Beth Stevens and Pascal Kaeser for Dissertation Advisory Committee guidance. Thanks to the Young-Pearse lab members for their valuable advice with experimental design and feedback on the manuscript. This study was supported by Edward R. and Anne G. Lefler Center Predoctoral Fellowship (G.T.), Larry L. Hillblom Foundation nos. 2025-A-175-FEL (J.P.C.), NIH grant U01AG072572 (P.L.D.J.), Chan-Zuckerberg Initiative grant CS-02018-191971 (P.L.D.J.), NIA grant R01AG055909 (T.L.Y.-P.), and Alzheimer’s Association Zenith Award (T.L.Y.-P.). ROSMAP is supported by P30AG10161, P30AG72975, R01AG17917, R01AG015819, U01AG072572, U01AG046152, and U54AG09066.

## Author contributions

Conceptualization, G.T. and T.L.Y.-P.; methodology, G.T., E.S.K., S.E.H., J.P.C., V.C.H., H.D., G.M.F., A.J.C., N.A., J.K.S., A.C., C.R.M., D.M.D., N.T.S., P.L.D.J., M.B.-J., T.L.Y.-P.; software, G.T., E.S.K, S.E.H., F.A., C.R.B.; formal analysis, G.T., E.S.K., S.E.H., F.A., C.R.B. D.M.D. and T.L.Y.-P.; investigation, G.T., E.S.K., S.E.H.; resources, D.A.B, N.T.S, P.L.D.J and T.L.Y.-P.; data curation, G.T., E.S.K., S.E.H and T.L.Y.-P.; writing, G.T. and T.L.Y.-P.; visualization, G.T. and T.L.Y.-P.; supervision, T.L.Y.-P.; project administration, T.L.Y.-P.; funding acquisition G.T., D.A.B, P.L.D.J., M.B.-J., and T.L.Y.-P.

---

**Extended Data Fig. 1:**
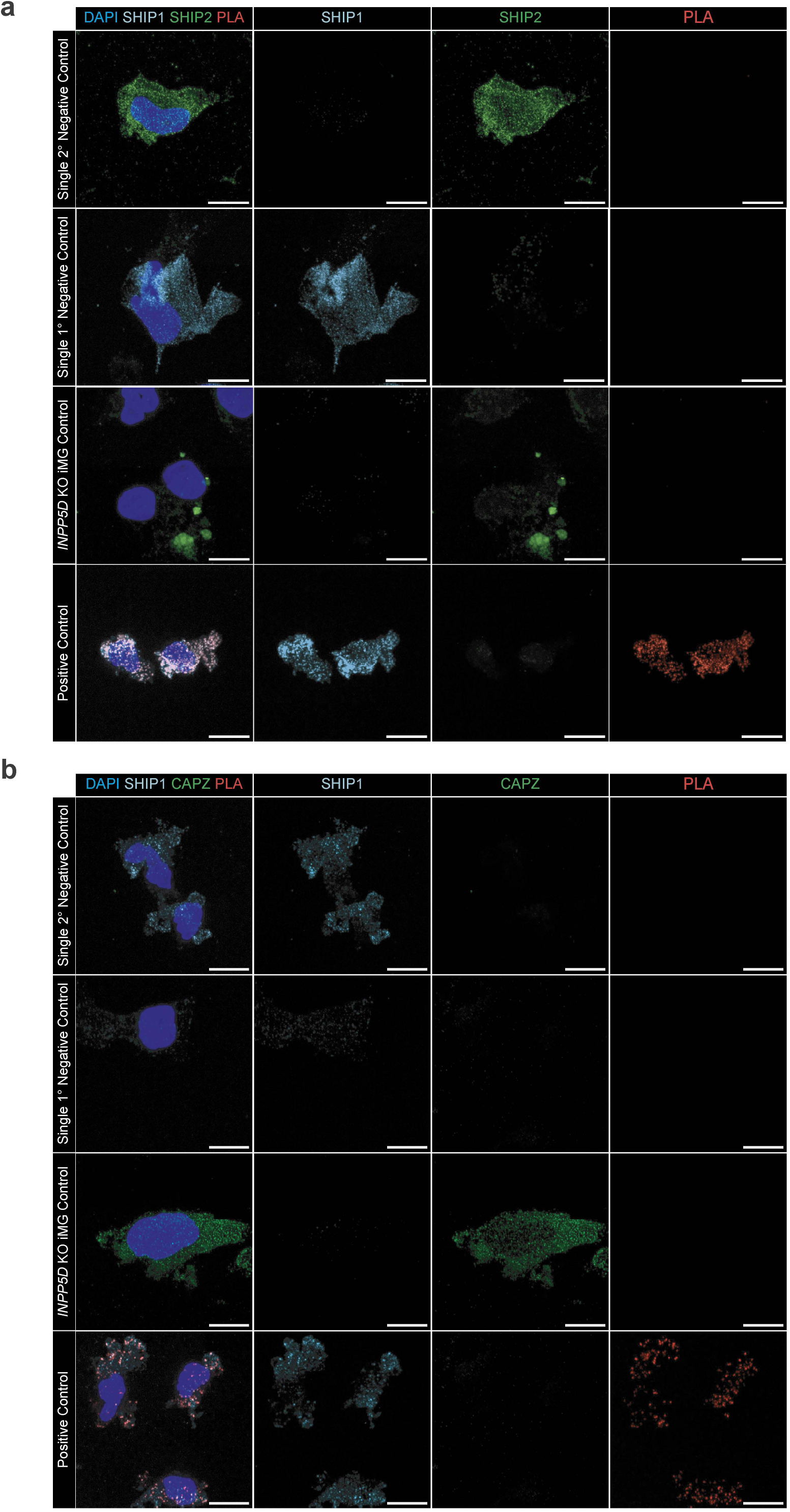
Controls for the proximity ligation assay (PLA) Representative confocal images of positive and negative controls for proximity ligation assays in (a) Fig. 1g and (b) Fig. 1h. PLA relies on a pair of secondary antibodies conjugated with oligonucleotides, which, when within 40 nm, initiate rolling circle amplification, resulting in a highly sensitive fluorescent signal at single-molecule resolution^82^. Negative controls, which either omitted the primary or secondary antibodies or used *INPP5D* KO iMGs, yielded no detectable signal. In contrast, the positive control, using both PLA probes directed against the SHIP1 primary antibody’s host species (rabbit), produced widespread PLA signal. Scale bars: 10 μm.

**Extended Data Fig. 2:**
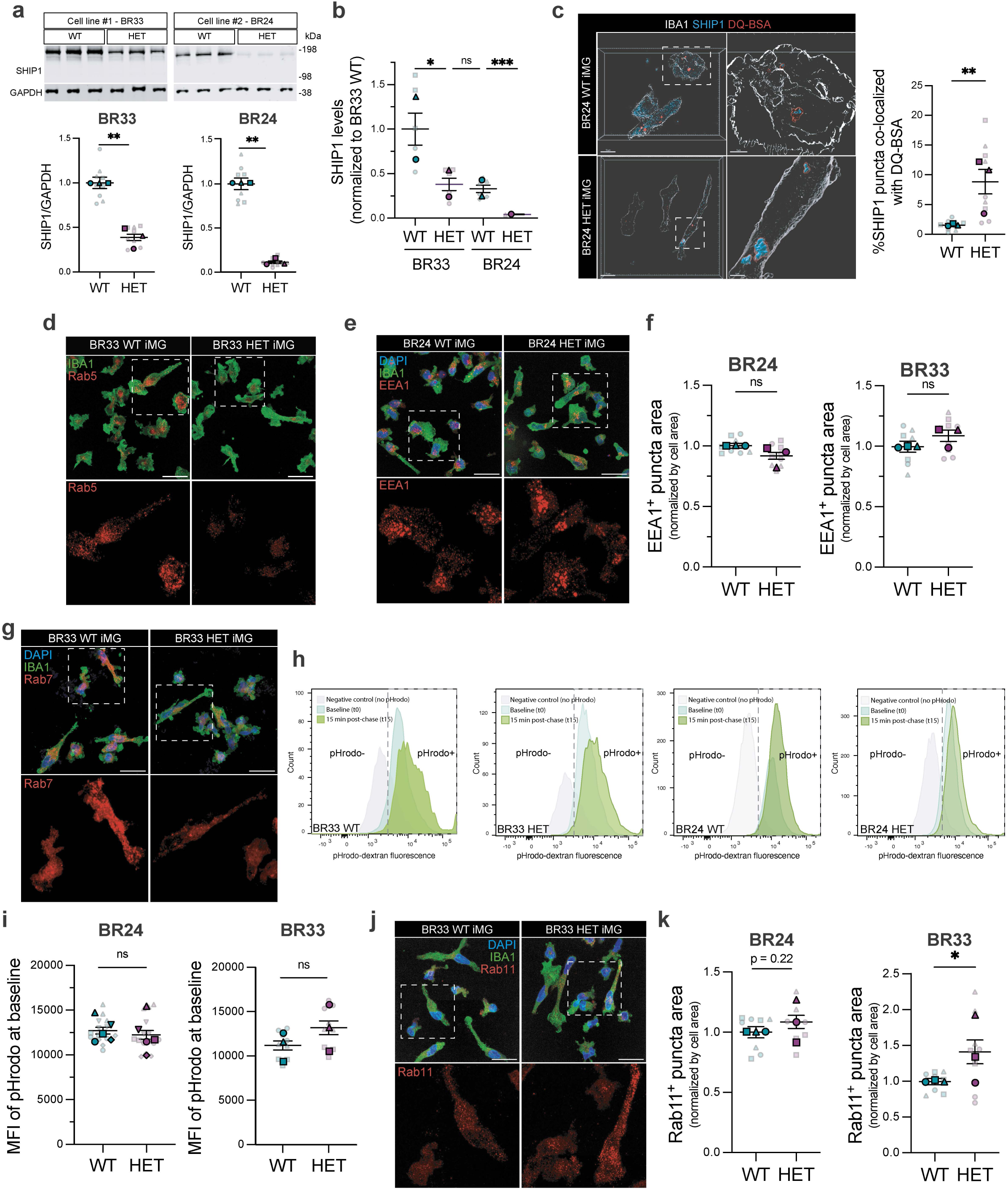
Additional characterization of endosomal and degradative phenotypes in SHIP1-deficient human microglia. a. Representative western blots and quantification of SHIP1 protein abundance in WT and *INPP5D* haploinsufficient (HET) iMGs from two independent genetic backgrounds, BR33 and BR24. b. Quantification of SHIP1 protein abundance normalized to BR33 WT iMGs across both genetic backgrounds. One-way ANOVA with Tukey’s multiple comparisons analysis test was performed using averages from each differentiation. c. Representative 3D reconstructed image of IBA1+ BR33 iMGs immunostained for SHIP1 after 1-hour incubation and 2-hour chase of DQ-Red BSA and quantification of the percentage of SHIP1 puncta co-localized with DQ-BSA puncta in WT vs HET iMGs. The insets show the co-localized SHIP1-DQ-BSA puncta. See Supplementary Videos 1 and 2 for the full images. d. Representative confocal images of Rab5 in BR33 iMGs. Scale bars: 25 μm. e. Representative confocal images of EEA1 in BR24 iMGs. Scale bars: 25 μm. f. Quantification of EEA1-positive puncta area normalized to cell area in BR24 and BR33 WT and HET iMGs. g. Representative confocal images of Rab7 in BR33 iMGs. Scale bars: 25 μm. h. Representative histograms and gating strategy for BR33 and BR24 WT and HET iMGs treated with pHrodo-dextran for 5 minutes and measured either directly after the 5-minute treatment (baseline, t0) or after the pHrodo signal is chased for 15 minutes (t15). i. Quantification of baseline (t0) pHrodo-dextran fluorescence intensity in BR24 and BR33 WT and HET iMGs. j. Representative confocal images of Rab11 in BR33 iMGs. Scale bars: 25 μm. k. Quantification of Rab11-positive puncta area normalized to cell area in BR24 and BR33 WT and HET iMGs. Unless otherwise stated, data is shown as mean ± SEM, normalized to WT conditions, obtained from n=3 independent differentiations with 3 technical replicates per differentiation. Mixed-effects analysis was performed with genotype and differentiation as fixed factors. The reported p-value corresponds to the main effect of genotype, averaged across differentiations. ****p < 0.0001, ***p < 0.001, **p < 0.01, *p<0.05; ns, not significant. For all scatter plots, different shapes delineate data from separate differentiations, average values per differentiation are shown, with faded dots representing technical replicates (wells) within each differentiation.

**Extended Data Fig. 3.**
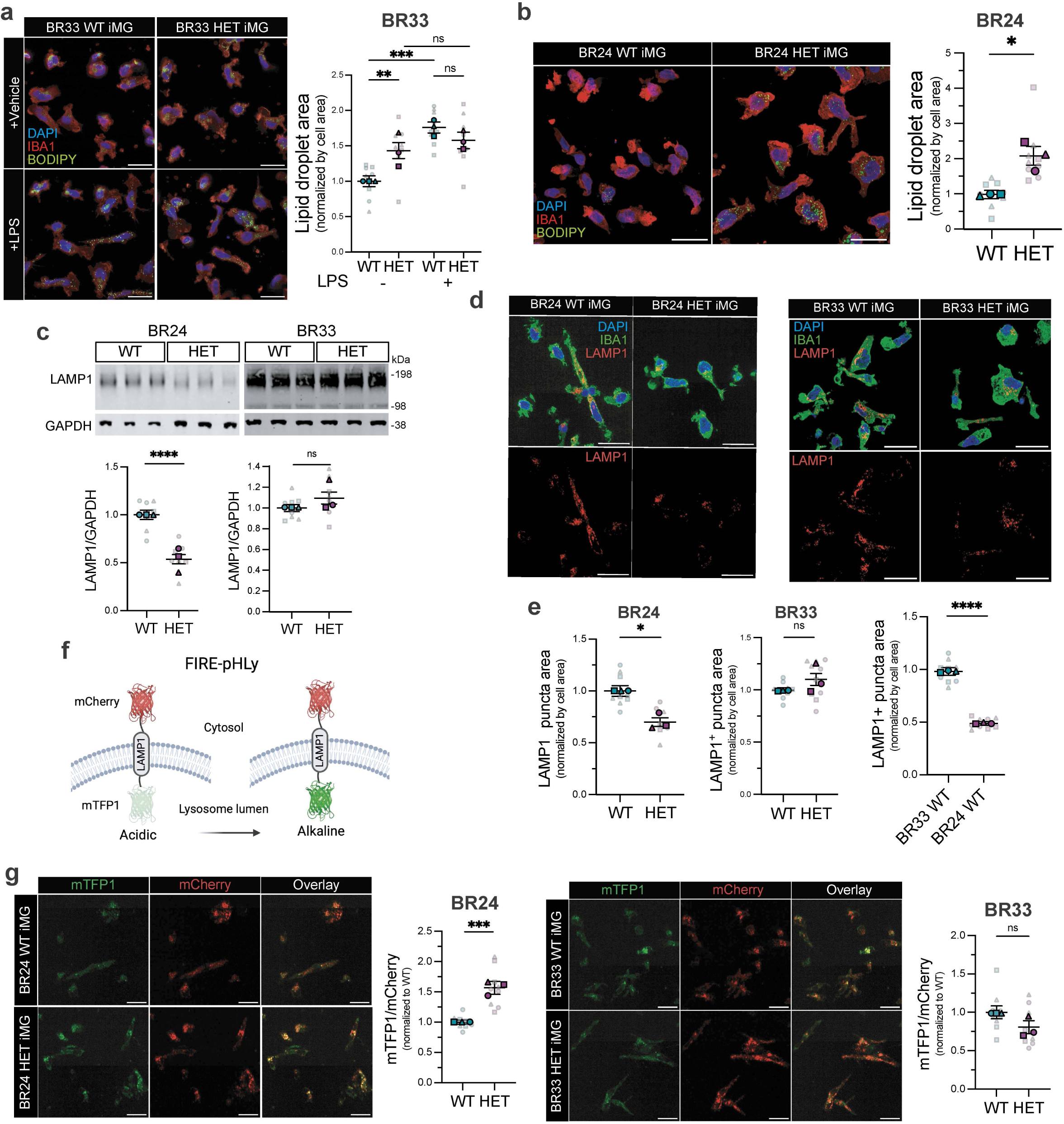
Conserved lipid droplet accumulation and variable lysosomal phenotypes in SHIP1-deficient human microglia. a. Representative confocal images and quantification of BODIPY 493/503 with vehicle or 6-hour LPS treatment in BR33 iMGs. Scale bars: 25 μm. b. Representative confocal images and quantification of BODIPY 493/503 in BR24 iMGs. Scale bars: 25 μm. c. Representative Western blot and quantification of LAMP1, normalized by GAPDH, in BR33 and BR24 WT vs HET iMGs. d. Representative confocal images of LAMP1 in BR24 and BR33 iMGs. Scale bars: 25 μm. e. Quantification of LAMP1+ puncta area, normalized by cell area, in BR24 and BR33 iMGs. f. Schematic of FIRE-pHLy lysosomal pH biosensor. mTFP1, which faces the lysosomal lumen, increases in fluorescence in alkaline vs acidic pH. mCherry, which faces the cytosol, is constitutively fluorescent. Created with BioRender.com. g. Representative images for mTFP1 and mCherry and quantification of mTFP1/mCherry ratio in BR24 and BR33 iMGs. Scale bars: 25 μm. Unless otherwise stated, data is shown as mean ± SEM, normalized to WT conditions, obtained from n=3 independent differentiations with 3 technical replicates per differentiation. Mixed-effects analysis was performed with genotype and differentiation as fixed factors. The reported p-value corresponds to the main effect of genotype, averaged across differentiations. ****p < 0.0001, ***p < 0.001, **p < 0.01, *p<0.05; ns, not significant. For all scatter plots, different shapes delineate data from separate differentiations, average values per differentiation are shown, with faded dots representing technical replicates (wells) within each differentiation.

**Extended Data Fig. 4.**
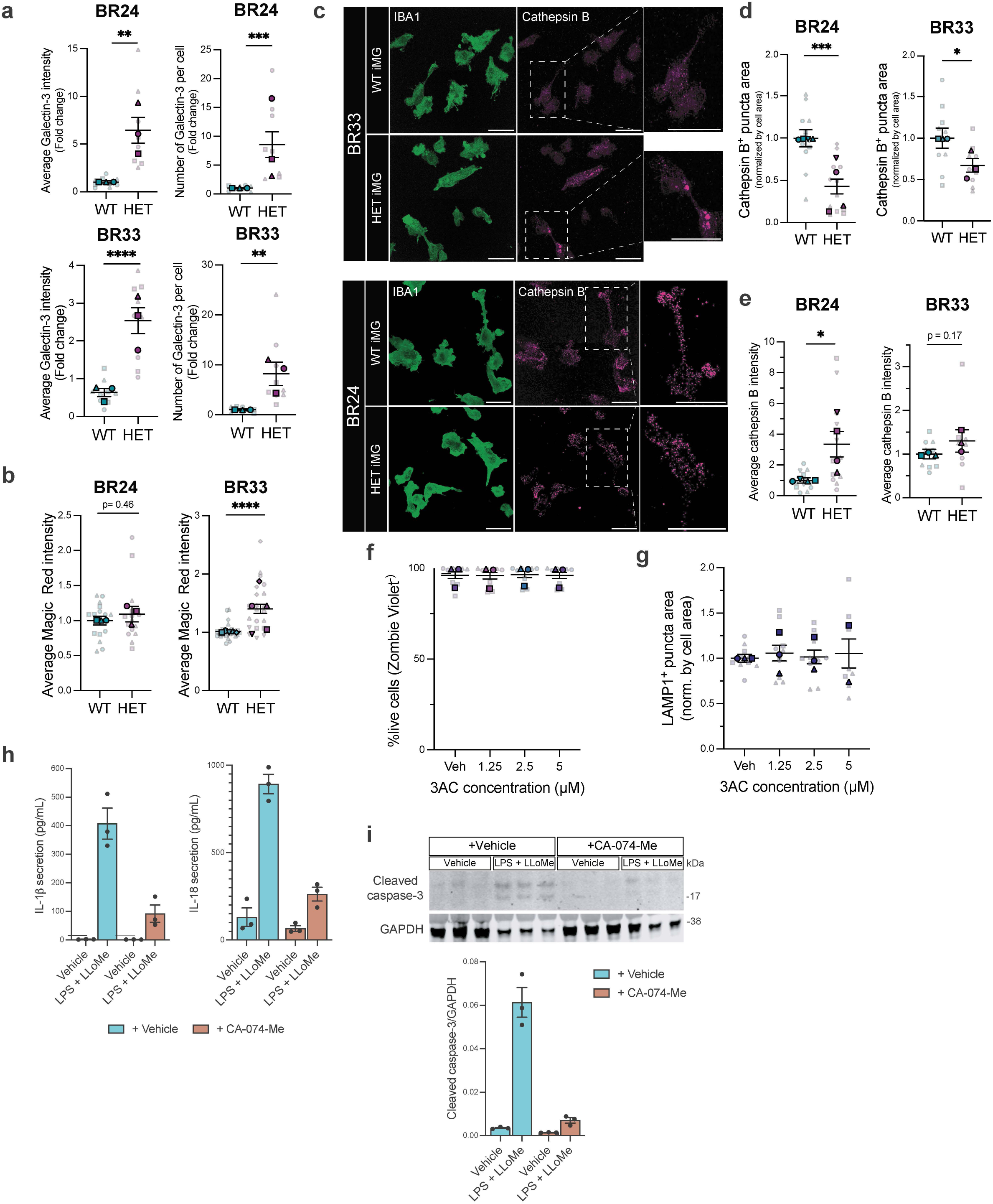
SHIP1 deficiency promotes lysosomal membrane damage and cathepsin B-dependent inflammasome activation. a. Average galectin-3 signal intensity and galectin-3+ puncta per cell in BR24 and BR33 WT and HET iMGs. b. Quantification of Magic Red intensity (cathepsin B activity) in BR24 and BR33 WT and HET iMGs. c. Representative confocal images of BR33 and BR24 WT and HET iMGs immunostained for cathepsin B, IBA1 and DAPI. Scale bars: 10 μm. d. Quantification of cathepsin B+ puncta area, normalized by IBA1 area in BR24 and BR33 WT and HET iMGs. e. Quantification of average cathepsin B intensity in BR24 and BR33 WT and HET iMGs. f. Quantification of cell viability following treatment with increasing concentrations of 3AC. g. Quantification of LAMP1+ puncta area following treatment with increasing concentrations of 3AC. h. IL-18 and IL-1β secretion in WT iMGs treated with vehicle (water) LPS for 4 hours, followed by LLoMe for 2 hours, with 1-hour pre-treatment with CA-074-Me or vehicle (DMSO). i. Representative Western blot and quantification of cleaved caspase-3 levels, normalized by GAPDH, in iMGs with vehicle or LPS + LLoMe treatment and vehicle or CA-074-Me pre-treatment. Unless otherwise stated, data is shown as mean ± SEM, normalized to WT conditions, obtained from n=3 independent differentiations with 3 technical replicates per differentiation. Mixed-effects analysis was performed with genotype and differentiation as fixed factors. The reported p-value corresponds to the main effect of genotype, averaged across differentiations. ****p < 0.0001, ***p < 0.001, **p < 0.01, *p<0.05; ns, not significant. For all scatter plots, different shapes delineate data from separate differentiations, average values per differentiation are shown, with faded dots representing technical replicates (wells) within each differentiation.

**Extended Data Fig. 5:**
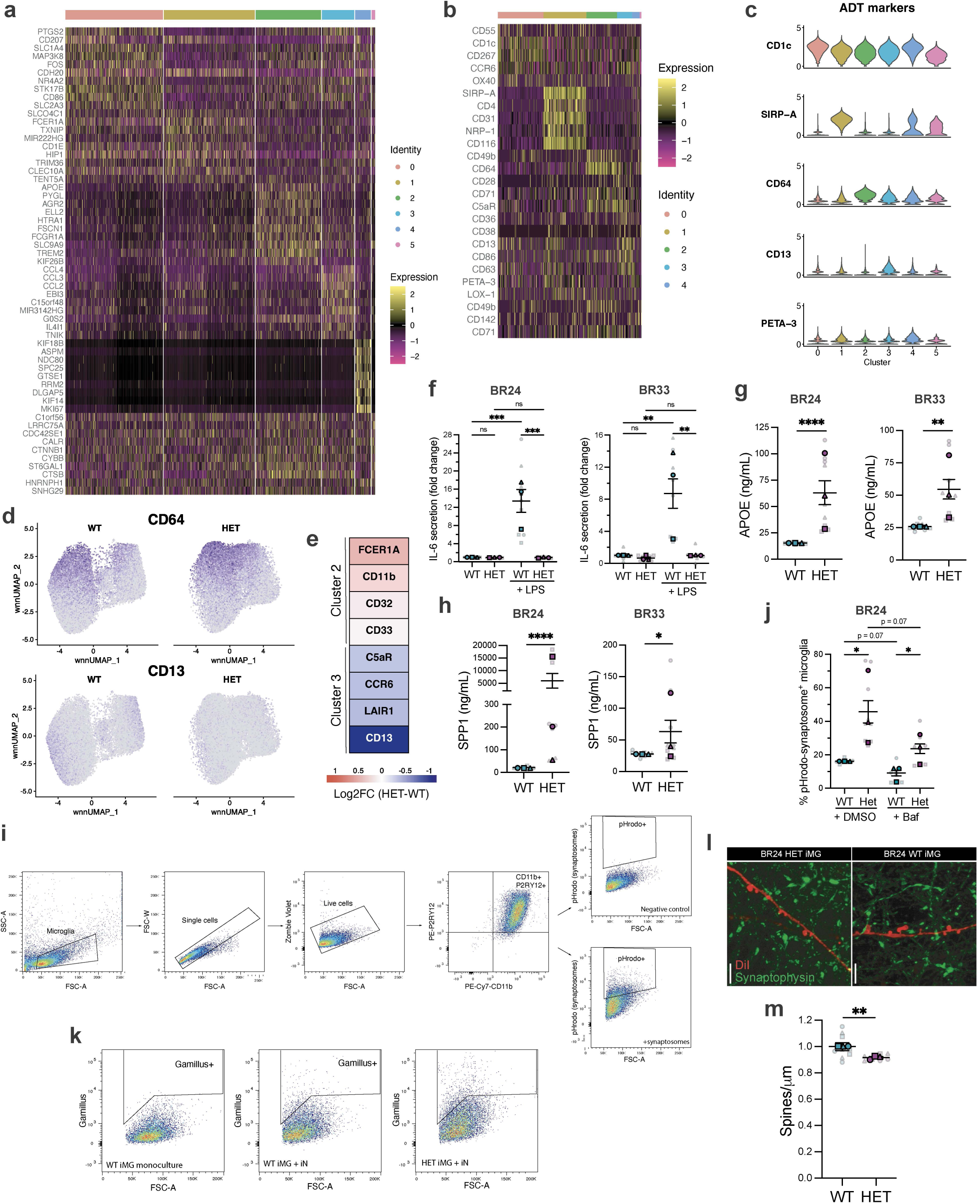
***INPP5D* haploinsufficiency induces a shift in microglial states** a. Heatmap of the top gene markers of each cluster. See Supplementary table 4 for the full list of markers. b. Heatmap of top ADT markers in each cluster. See Supplementary Table 6 for the full list of ADT markers. c. Violin plots of top ADT markers for clusters 1-4. d. Feature plots showing the expression of CD64 and CD13 across all iMGs in UMAP space. e. Heatmap showing log2 fold-change (log2FC) in expression of selected cluster 2 and cluster 3 differentially expressed proteins across WT and HET iMGs within the respective clusters. Red indicates proteins upregulated in HET iMGs, and blue indicates proteins downregulated in HET iMGs relative to WT. See Supplementary table 7 for differential expression of proteins across WT and HET iMGs within each cluster. f. IL-6 secretion, measured via ELISA (MSD) from BR24 and BR33 WT and HET iMGs treated with LPS or vehicle (water) for 6 hours. g. APOE secretion, measured via ELISA (MSD) from BR24 and BR33 WT and HET iMGs. h. SPP1 secretion, measured via ELISA (MSD) from BR24 and BR33 WT and HET iMGs. i. Flow cytometry gating strategy to measure pHrodo fluorescence in live P2RY12+ CD11b+ iMGs. j. Percentage of BR33 WT and HET iMGs positive for pHrodo-labeled synaptosomes in the presence or absence of bafilomycin A1 (Baf), measured via flow cytometry. Two-way ANOVA was performed using averages from each differentiation. k. Flow cytometry gating strategy to measure Gamillus intensity in iMG monocultures (negative control) and co-cultures. l. Representative confocal images of dendrites (stained with DiI) of iNs co-cultured with BR24 WT or HET iMGs for 3 days. Scale bars: 5 μm. m. Quantification of spine density in iNs co-cultured with BR24 WT or HET iMGs. Unless otherwise stated, data is shown as mean ± SEM, normalized to WT conditions, obtained from n=3 independent differentiations with 3 technical replicates per differentiation. Mixed-effects analysis was performed with genotype and differentiation as fixed factors. The reported p-value corresponds to the main effect of genotype, averaged across differentiations. ****p < 0.0001, ***p < 0.001, **p < 0.01, *p<0.05; ns, not significant. For all scatter plots, different shapes delineate data from separate differentiations, average values per differentiation are shown, with faded dots representing technical replicates (wells) within each differentiation.

**Extended Data Fig. 6:**
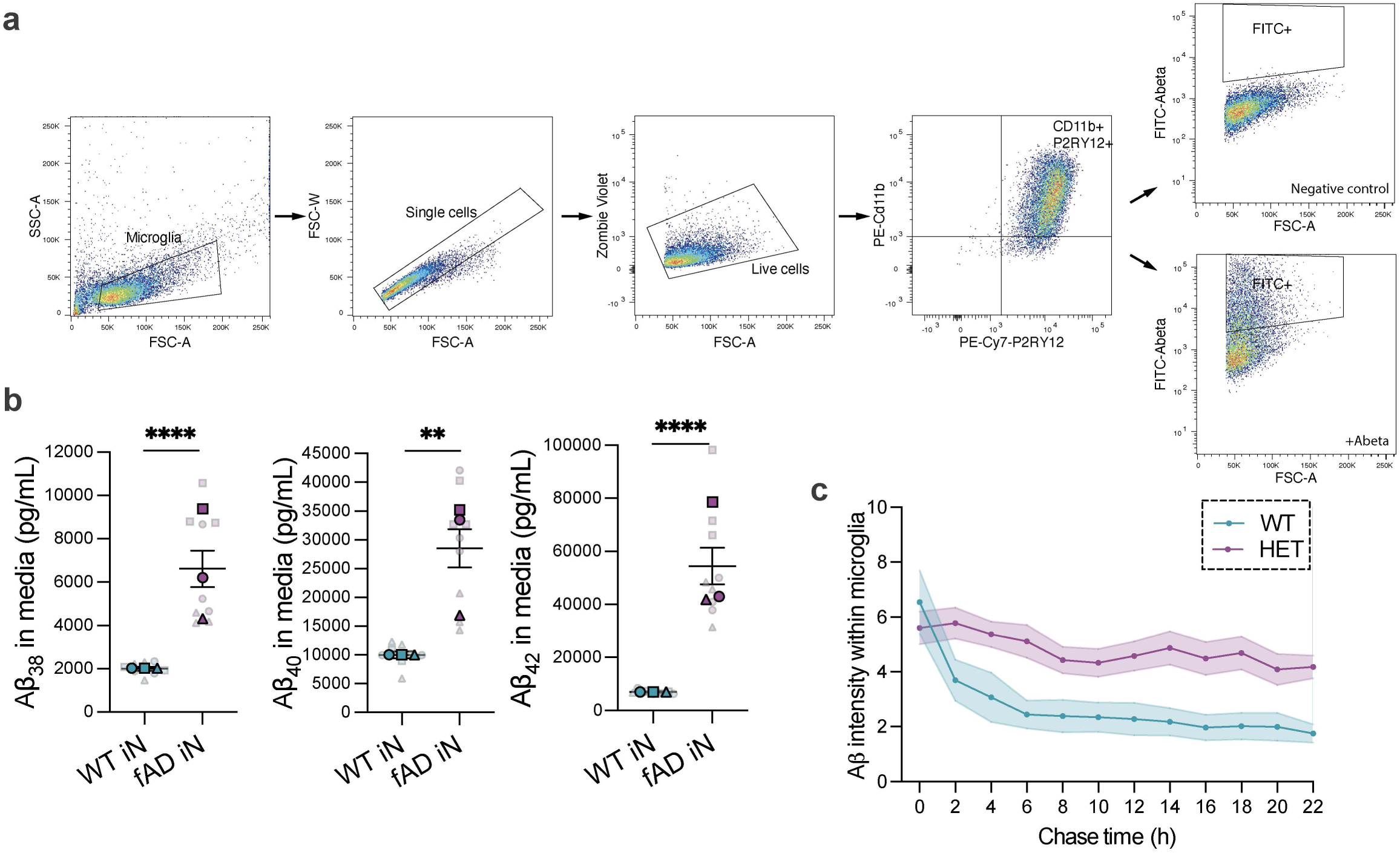
Measurements of Aβ uptake and degradation in iMGs. a. Flow cytometry gating strategy to measure FITC-Abeta fluorescence in live P2RY12^+^ CD11b^+^ iMGs. b. ELISA measurements of secreted Aβ_38_, Aβ_40_ and Aβ_42_ from WT and fAD iN monocultures. Quantification shown for n=3 differentiations with 3 technical replicates per differentiation. Mixed-effects analysis was performed with genotype and differentiation as fixed factors. The reported p-value corresponds to the main effect of genotype, averaged across differentiations. c. IncuCyte quantification of FITC-Aβ intensity within microglia over time. Quantification shown for n=1 differentiation with 7 technical replicates (wells), averaged from 9 images per well.

**Extended Data Fig. 7:**
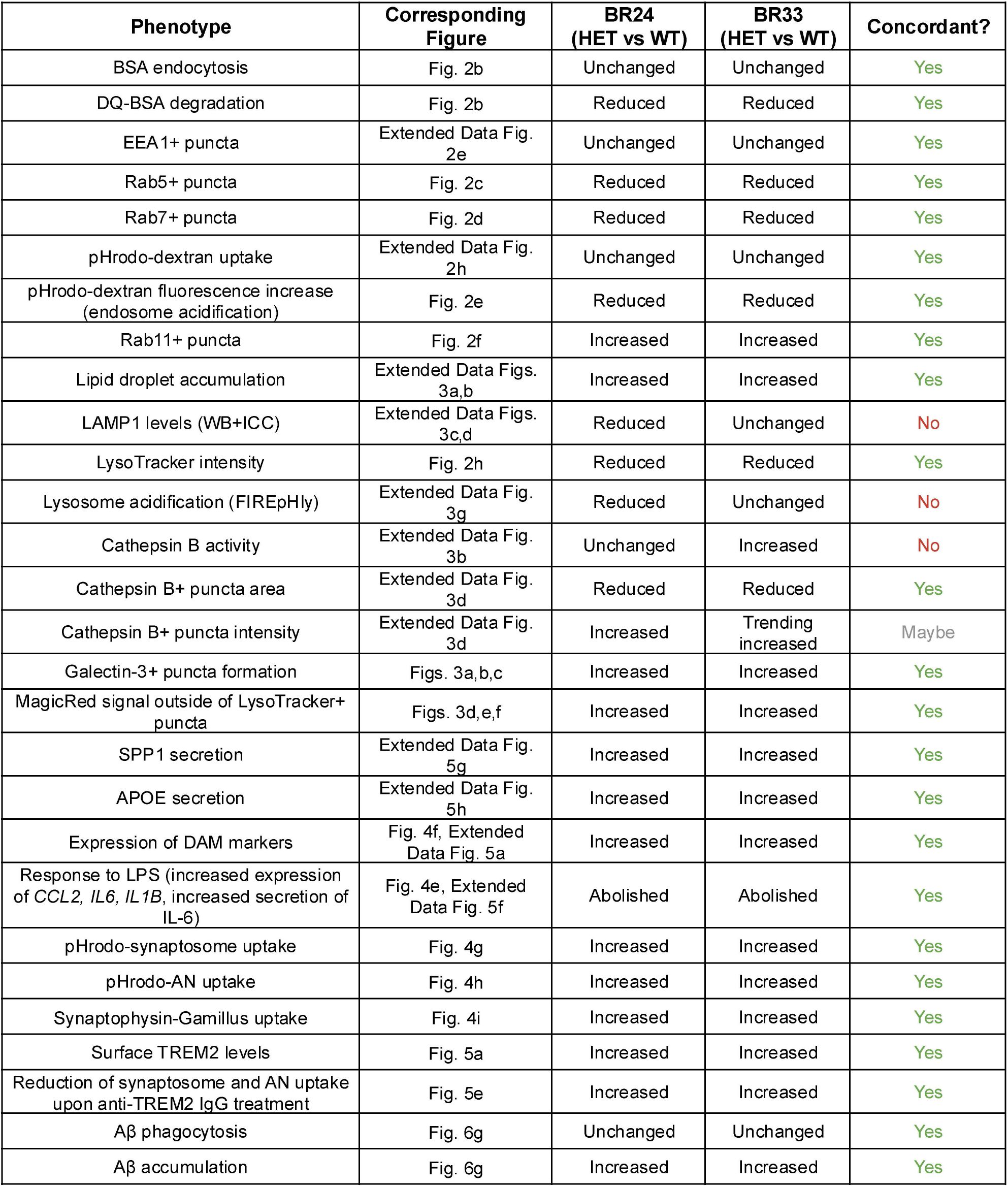
Concordance of *INPP5D* haploinsufficiency phenotypes across two independent iMG genetic backgrounds. Summary of phenotypes observed in *INPP5D* haploinsufficient (HET) iMGs across BR24 and BR33 genetic backgrounds. Phenotypes were classified as concordant when both genetic backgrounds exhibited significant changes in the same direction relative to WT iMGs.

**Extended Data Fig. 8:**
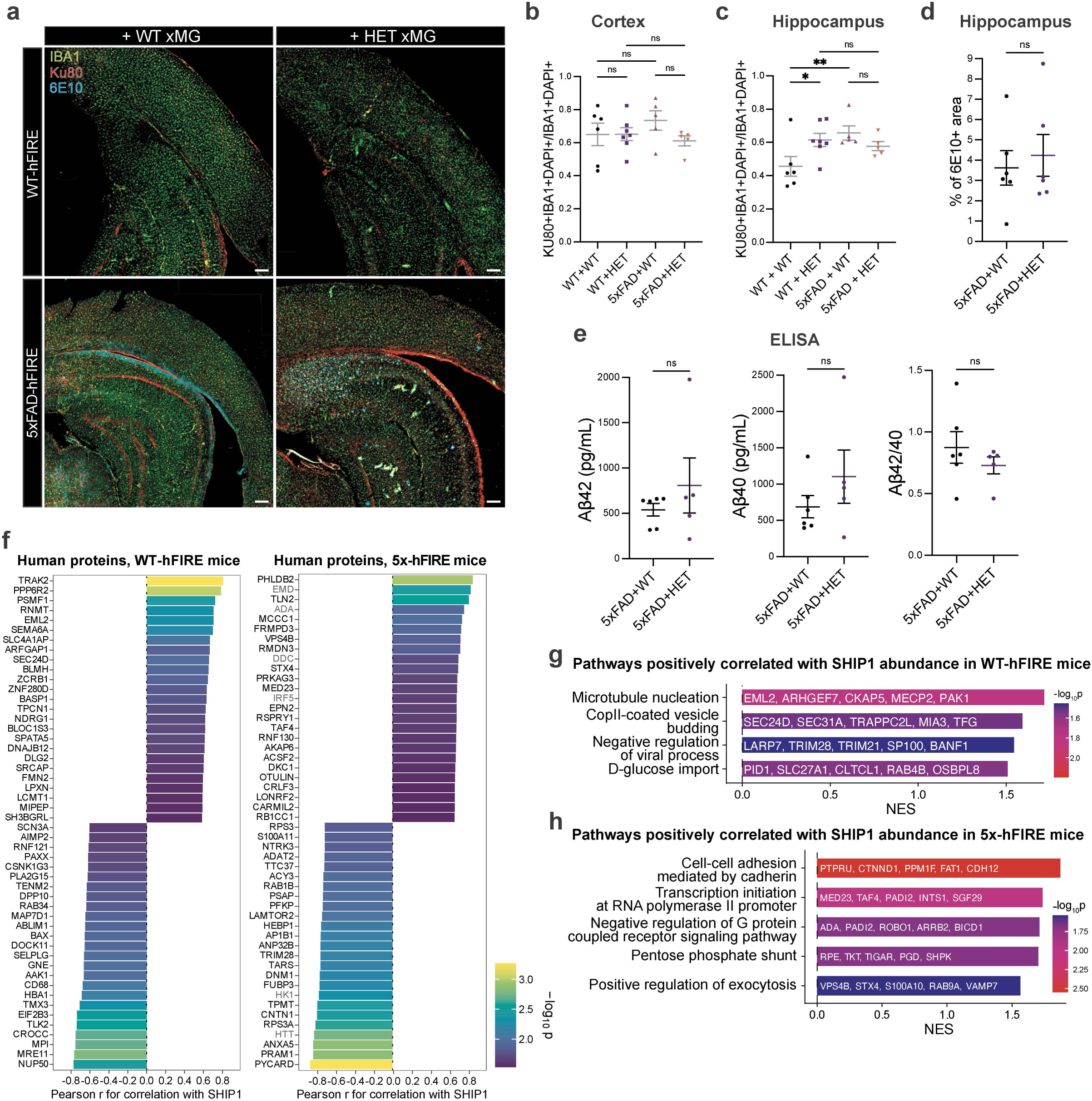
Additional characterization of xenografted human xMGs and SHIP1-associated proteomic programs in WT-hFIRE and 5x-hFIRE brains. a. Representative low-magnification images of WT and *INPP5D* haploinsufficient (HET) human xMG engraftment in WT-hFIRE and 5x-hFIRE brains. Human xMGs were identified as Ku80+, IBA1+ cells. 6E10 labels amyloid plaques. Scale bars: 100 μm. b. Ratio of IBA1+Ku80+DAPI+ microglia to IBA1+DAPI+ in cortex across experimental groups. c. Ratio of IBA1+Ku80+DAPI+ microglia to IBA1+DAPI+ in hippocampus across experimental groups. d. Quantification of plaque burden in hippocampus measured as percentage of 6E10+ area in xenografted 5x-hFIRE brains. e. ELISA quantification of Aβ42, Aβ40, and Aβ42/40 ratio in WT and HET xMG-engrafted 5x-hFIRE brains. f. Human proteins most positively and negatively correlated with SHIP1 abundance in WT-hFIRE and 5x-hFIRE brains. See Supplementary Tables 14 and 15 for correlation of all proteins with SHIP1 in WT-hFIRE and 5x-hFIRE brains. g. Selected pathways positively correlated with SHIP1 abundance in human proteins derived from WT-hFIRE-engrafted brains. See Supplementary Table 16 all GO GSEA results. h. Selected pathways positively correlated with SHIP1 abundance in human proteins derived from 5x-hFIRE-engrafted brains. See Supplementary Table 17 all GO GSEA results.

**Extended Data Fig. 9:**
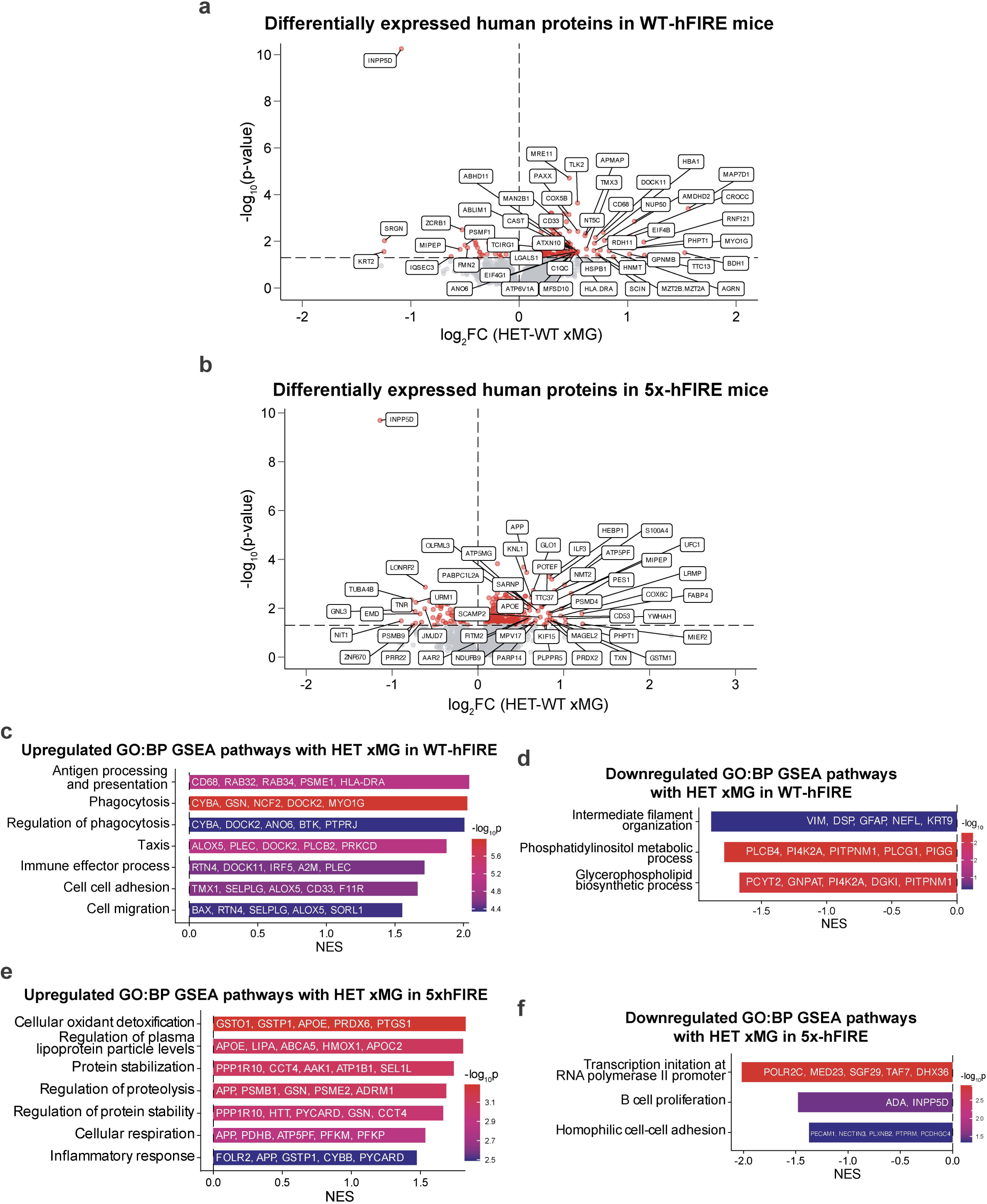
Differential proteomic changes associated with SHIP1 deficiency in xenografted human microglia. a. Volcano plot of differentially abundant human proteins across HET and WT xMGs engrafted into WT-hFIRE brains. Dashed line indicates p-value significance threshold. See Supplementary Table 18 for all differentially abundant human proteins in WT-FIRE mice. b. Volcano plot of differentially expressed human proteins comparing HET and WT xMGs engrafted into 5xFAD-hFIRE brains. Dashed line indicates p-value significance threshold. See Supplementary Table 19 for all differentially abundant human proteins in 5xFAD-hFIRE mice. c. Selected GO biological process pathways positively enriched in HET xMGs relative to WT xMGs in WT-hFIRE brains. See Supplementary Table 20 all GO GSEA results. d. Selected GO biological process pathways negatively enriched in HET xMGs relative to WT xMGs in WT-hFIRE brains. See Supplementary Table 20 all GO GSEA results. e. Selected GO biological process pathways positively enriched in HET xMGs relative to WT xMGs in 5xFAD-hFIRE brains. See Supplementary Table 21 all GO GSEA results. f. Selected GO biological process pathways negatively enriched in HET xMGs relative to WT xMGs in 5xFAD-hFIRE brains. See Supplementary Table 21 all GO GSEA results.

