## Supplementary material for "INPP5D/SHIP1 regulates endo-lysosomal function and cargo-selective phagocytosis in human microglia": Description of supplementary files

**Supplementary Video 1:** 3D reconstruction of SHIP1 and DQ-BSA signal in WT iMGs

**Supplementary Video 2:** 3D reconstruction of SHIP1 and DQ-BSA signal in HET iMGs

**Supplementary Table 1:** SHIP1 binding partners identified via IP-MS

**Supplementary Table 2:** GO cellular component terms associated with SHIP1 binding partners in iMGs identified via IP-MS

**Supplementary Table 3:** Reactome pathways associated with SHIP1 binding partners in iMGs identified via IP-MS

**Supplementary Table 4:** RNA markers of microglia clusters in the CITE-seq dataset identified via the R package FindAllMarkers

**Supplementary Table 5:** Surface protein markers of microglia clusters in the CITE-seq dataset identified via the R package FindAllMarkers

**Supplementary Table 6:** GO enrichment results for the RNA markers in each cluster in the CITE-seq dataset

**Supplementary Table 7:** Differentially expressed proteins across WT and HET iMGs within each cluster in the CITE-seq dataset

**Supplementary Table 8:** Metadata for ROSMAP donor cell lines 8.used for iMG generation

**Supplementary Table 9:** Proteins correlated with SHIP1 abundance across genetically diverse iMGs

**Supplementary Table 10:** KEGG GSEA of proteins correlated with SHIP1 abundance across genetically diverse iMGs

**Supplementary Table 11:** GO biological process GSEA of proteins correlated with SHIP1 abundance across genetically diverse iMGs

**Supplementary Table 12:** Differential protein abundance associated with rs10933431-G allele dosage

**Supplementary Table 13:** GO biological process GSEA of proteins associated with rs10933431-G allele dosage

**Supplementary Table 14:** Human proteins correlated with SHIP1 abundance in xMG-engrafted WT-hFIRE mice

**Supplementary Table 15:** Human proteins correlated with SHIP1 abundance in xMG-engrafted 5xFAD-hFIRE mice

**Supplementary Table 16:** GO biological process GSEA of proteins correlated with SHIP1 abundance in xMG-engrafted WT-hFIRE mice

**Supplementary Table 17:** GO biological process GSEA of proteins correlated with SHIP1 abundance in xMG-engrafted 5xFAD-hFIRE mice

**Supplementary Table 18:** Differential protein abundance analysis results across HET and WT XMGs engrafted into WT-hFIRE brains

**Supplementary Table 19:** Differential protein abundance analysis results across HET and WT xMGs engrafted into 5xFAD-hFIRE brains

**Supplementary Table 20:** GO biological process GSEA of differentially abundant proteins across HET and WT xMGs engrafted into WT-hFIRE brains

**Supplementary Table 21:** GO biological process GSEA of differentially abundant proteins across HET and WT xMGs engrafted into 5xFAD-hFIRE brains
